# Selection and structural characterisation of anti-TREM2 scFvs that reduce levels of shed ectodomain

**DOI:** 10.1101/2021.02.07.430123

**Authors:** Aleksandra Szykowska, Yu Chen, Thomas B Smith, Charlotta Preger, Jingjing Yang, Dongming Qian, Shubhashish M Mukhopadhyay, Edvard Wigren, Stephen J Neame, Susanne Gräslund, Helena Persson, Peter J Atkinson, Elena Di Daniel, Emma Mead, John Wang, John B Davis, Nicola A Burgess-Brown, Alex N Bullock

## Abstract

Single point mutations in TREM2, a receptor expressed by microglia in the brain, are associated with an increased risk of neurodegeneration including Alzheimer’s disease. Numerous studies support a role for TREM2 in sensing damaging stimuli and triggering signalling cascades necessary for neuroprotection. Despite its significant role, ligands and regulators of TREM2 activation, and the mechanisms governing TREM2-dependent responses and its cleavage from the membrane, remain poorly characterised. Here, we present phage display generated scFv antibody binders to human TREM2 ectodomain. Cocrystal structures revealed the binding of two scFvs to an epitope on the globular TREM2 domain distal to the putative ligand-binding site. Enhanced functional activity was observed for oligomeric scFv species which inhibited the production of soluble TREM2 in a HEK293 cell model. We hope that detailed characterisation of their epitopes and properties will facilitate the use of these renewable binders as structural and functional biology tools for TREM2 research.

## Introduction

Late-onset Alzheimer’s disease (LOAD) is the most common type of dementia, characterised by accumulation of extracellular amyloid-β (Aβ) aggregates and intracellular neurofibrillary tangles of hyper-phosphorylated tau, with a long prodromal phase followed by cognitive decline. Despite the urgency and necessity to develop therapeutics, there are currently no approved drugs which cure or slow the underlying progression of Alzheimer’s disease (AD) and patients are still reliant on symptomatic treatments discovered in the mid-last century (Long and Holtzman, 2019).

After some late-stage clinical trials targeting Aβ failed to meet their desired endpoints and several large genome-wide association studies (GWAS) linked genes coding for components of the immune response to AD, neuro-inflammation has become an area of intense research for therapeutics (Block *et al*., 2007; Lambert *et al*., 2013). Among the genes identified by GWAS is *TREM2* (Triggering Receptor Expressed on Myeloid cells), which encodes a single transmembrane receptor expressed in myeloid-derived cells, including microglia in the central nervous system (CNS) (Guerreiro *et al*., 2013). Homozygous loss-of-function mutations in TREM2, or the associated adaptor protein DAP12, were previously identified to cause aggressive early-onset dementia in Nasu-Hakola disease (Paloneva *et al*., 2003). Since then, several point mutations in the extracellular domain of TREM2 have been linked to neurodegenerative disorders, highlighting the importance of TREM2 functions in brain health and homeostasis (Ulrich and Holtzman, 2017; Konishi and Kiyama, 2018). The most notable amino acid substitution, R47H, leads to four-fold increased risk of developing LOAD (Jonsson *et al*., 2013; Guerreiro *et al*., 2013). A stronger genetic effect is observed only in carriers of Apolipoprotein E (APOE) ε4, a potential TREM2 ligand, which has been implicated in TREM2 pathology (Krasemann *et al*., 2017; Shi and Holtzman, 2018; Parhizkar *et al*., 2019).

TREM2 has numerous putative functions including regulation of lipid and cholesterol metabolism, phagocytosis of myelin and Aβ, and generation of a microglial barrier around Aβ plaques (Yeh *et al*., 2016; Yuan *et al*., 2016; Jay *et al*., 2017; Ulland *et al*., 2017; Ulland and Colonna, 2018; Nugent *et al*., 2020). TREM2 has been reported to bind ligands as diverse as Aβ, lipids, myelin and lipoproteins (Wang *et al*., 2015; Yeh *et al*., 2016; Zhao *et al*., 2018). Recent findings have indicated that TREM2-dependent signalling is essential for the transcriptional definition of disease-associated microglia (DAM), a phenotype which is believed to be neuroprotective as it upregulates genes involved in phagocytosis, for example (Keren-Shaul *et al*., 2017).

TREM2 contains an immunoglobulin-like ectodomain followed by a flexible stalk region, a transmembrane domain and a short cytoplasmic tail. The stalk region can be cleaved by ADAM10/17 proteases to generate a soluble TREM2 fragment (sTREM2), while the C-terminal intramembranous domain is further cleaved by gamma-secretase (Wunderlich and Walter, 2013). sTREM2 can be detected in cerebrospinal fluid (CSF) and is increased in (Wang *et al*., 2020) patients with neuronal injury or CNS inflammatory diseases (Piccio *et al*., 2008; Rauchmann *et al*., 2019). Additionally, sTREM2 was found to be increased in patients at early symptomatic stages of AD and, crucially, correlated well with levels of phosphorylated tau in patients with tau pathology (Suárez-Calvet *et al*., 2016; Rauchmann *et al*., 2019). The importance of TREM2 cleavage in AD pathology is highlighted by the H157Y polymorphism at the protease cleavage site, which leads to excessive shedding of sTREM2 and increased risk of AD (Schlepckow *et al*., 2017; Thornton *et al*., 2017). It is unclear whether the increased risk is due to the resulting increased generation of additional sTREM2, which might be biologically active molecule (Zhong *et al*., 2019), alterations in intracellular signalling or functional properties of the remaining C-terminal fragments.

There is minimal published structural data for TREM2 ectodomain motifs responsible for ligand engagement or regulation of TREM2 functions (Kober *et al*., 2016; Sudom *et al*., 2018). While the ectodomain R47H mutant has been proposed to be defective in ligand binding, its crystal structure was solved only recently (Sudom *et al*., 2018). The authors concluded that the arginine substitution causes extensive remodelling in the neighbouring CDR2 loop of TREM2 with resulting loss of electron density. The same loop has been identified to interact with putative ligands in wild-type TREM2 crystals soaked with phosphatidylserine (Sudom *et al*., 2018). However, structures incorporating other ligands or the molecular mechanisms of TREM2 signalling remain to be fully elucidated.

Because of the potential therapeutic impact of targeting TREM2, we decided to generate single-chain variable antibody fragments (scFvs) against the human TREM2 ectodomain with which to study TREM2 structure and function. Agonist antibodies have already been reported in the recently published work (Schlepckow *et al*., 2020; Wang *et al*., 2020). The group of Schlepckow have shown that a full-length antibody specific to mouse TREM2 can decrease shedding and activate TREM2 signalling *in vitro*, and also lead to a significant reduction in amyloid plaques in 6-months old amyloid-beta precursor protein (APP) knock-in mice (Schlepckow *et al*., 2020). Antibodies might also be useful biochemical tools for studying the function of sTREM2 (Zhong *et al*., 2019).

There are currently no published structures of antibodies in complex with TREM2. Here we present our work on the generation and characterisation of four scFvs selected against the soluble ectodomain of human TREM2. The co-crystal structures of two of these antibody fragments in complex with TREM2 have been solved, revealing interactions with epitopes distal to the putative ligand binding site. Two scFvs reduced the shedding of sTREM2 from HEK293 cells.

Together, these data identify scFv antibody fragments as valuable reagents for cellular and structural studies of the TREM2 receptor and identify TREM2/ligand interfaces that may be used to modulate TREM2 function.

## Results

### Phage display selection of anti-TREM2 scFvs

To facilitate the development of renewable antibody binders against TREM2, we performed phage display selections using the SciLifeLib synthetic library of human scFvs similarly constructed and designed as previously reported (Säll *et al*., 2016; Preger *et al*., 2020). A TREM2 antigen comprising the ectodomain of human TREM2 (His19-Asp131) was expressed recombinantly from Sf9 insect cells and biotinylated on a C-terminal Avi tag for immobilisation on streptavidin-coated magnetic beads. Following four rounds of phage selections and initial binding analyses, four scFvs (scFv-1, 2, 3 and 4) were selected for further characterisation (**Figure 1**). All four scFv fragments were relatively poorly expressed when produced in *E. coli*. and, therefore, scFvs were prepared by secretion from insect cells, which allowed for higher yields, increased solubility and a reduction in endotoxin contamination (**Figures 2a-b**). The scFvs showed a tendency to dimerize as observed by size exclusion chromatography (SEC), as well as delayed elution from interaction with the Superdex medium **(Figure 2c),** which has previously been observed for nanobodies (Zimmermann *et al*., 2020).

**Figure 1.**
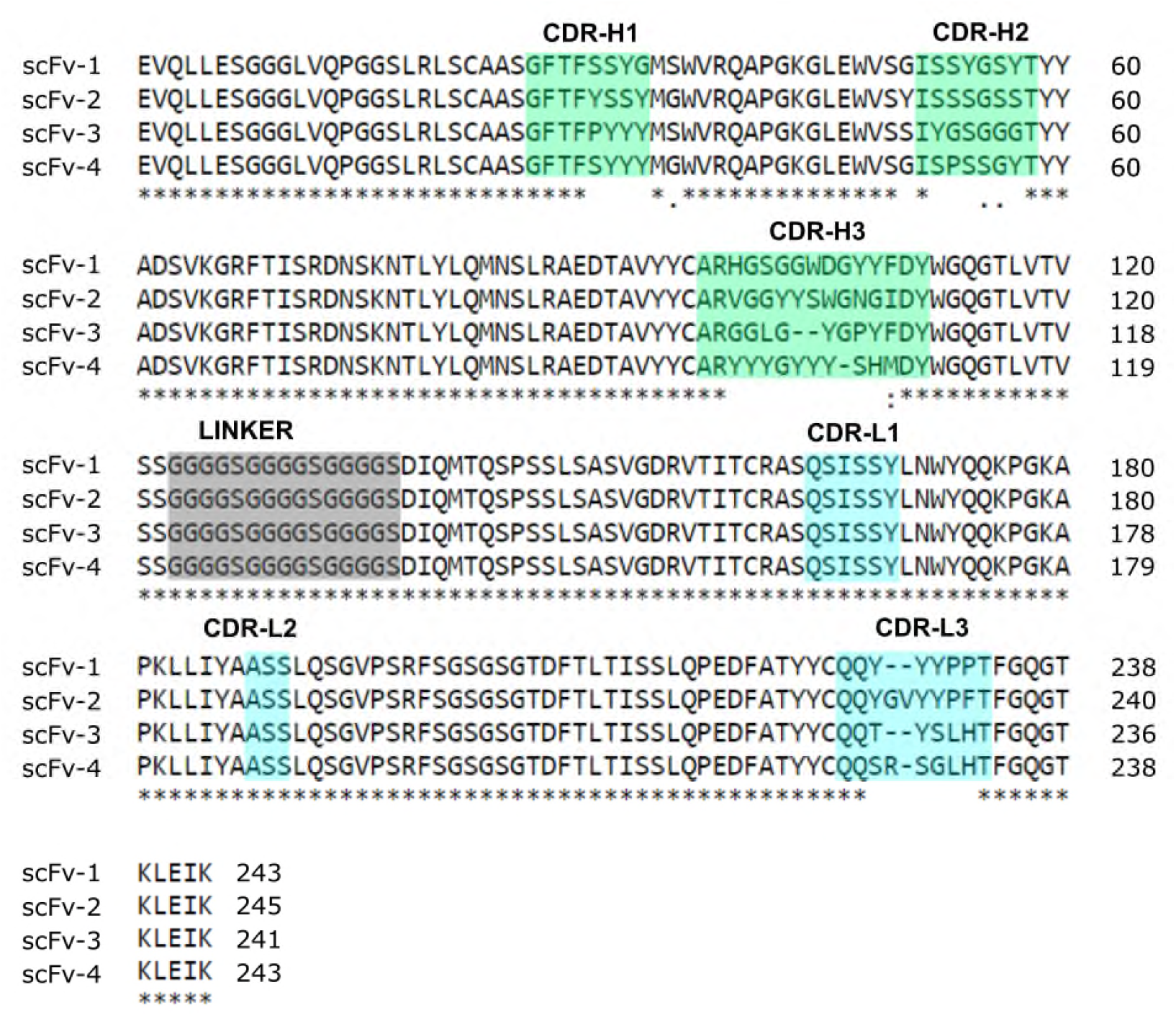
Multiple sequence alignment of anti-TREM2 scFv protein sequences. ScFv sequences are aligned by Clustal O with the three heavy chains CDR-H1-3 highlighted in green and light chains CDRL1-3 in blue, 15 a.a. linker in grey. Boundaries for CDRs are as defined by the IMGT nomenclature (Lefranc *et al*., 2003). The scFvs originate from the human synthetic phage library SciLifeLib (Preger *et al*., 2020), in which sequence diversity was introduced into four of the six CDRs: CDR-H1, CDR-H2, CDR-H3 and CDR-L3.

**Figure 2.**
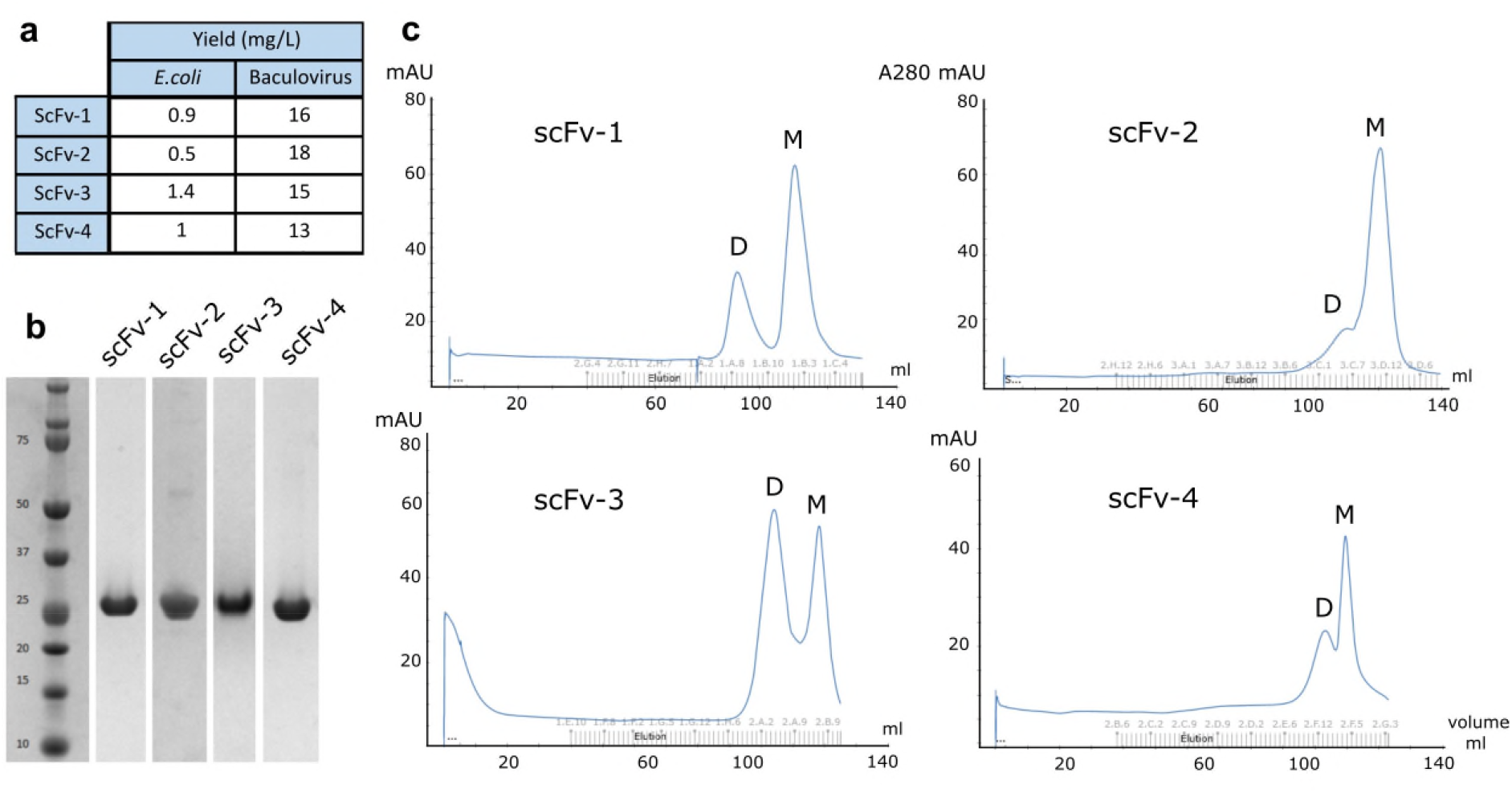
Optimisation of scFv production in a baculoviral expression system. (**a**) Expression yields of scFvs produced in *E. coli* TOP10 cells or Sf9 insect cells. (**b**) Coomassie blue-stained SDS-PAGE gel showing the final purity of scFvs after purification from Sf9 cells (image generated from different purification experiments). (**c**) Size exclusion chromatography of each scFv antibody fragment purified on a Superdex 200 16/60 column. Delayed elutions indicated scFv interactions with the Superdex media as well as a tendency for dimerization (D-dimer, M-monomer).

The binding of TREM2 to the scFvs, was characterised by surface plasmon resonance (SPR) to establish their relative binding affinities (**Figure S1**; apparent ^App^*K*¤ values are reported to account for potential avidity effects caused by scFv oligomerization (Preger *et al*., 2020). For these experiments, the human TREM2 antigen was cloned into a mammalian expression vector and produced as a longer biotinylated construct consisting of the full ectodomain and stalk region of TREM2 (His19-Ser174) that was then immobilised onto a streptavidin-coated sensor chip. Initial measurements on mixed oligomer samples revealed that all antibody fragments showed strong binding to TREM2 except for scFv-1, which proved to be a weak binder with an apparent equilibrium dissociation constant (^App^*K*_D_) in the micromolar range (data not shown). The binding kinetics of scFv-3 displayed moderate affinity (^App^*K*_D_ ~ 54 nM, **Figure S1b**). By contrast, scFv-2 and scFv-4 were characterised by high affinity binding, ^App^*K*_D_ values of ~ 1 nM (**Figures S1a and S1c**), although the accuracy of scFv-4 binding kinetics determination was limited by some non-specific interactions with the SPR sensor chip as well as the possible avidity effects due to scFv oligomerization **(Figure 2c).**

### Structure determination of TREM2-scFv antibody complexes

We hypothesised that scFv antibodies binding at different sites on the TREM2 ectodomain might differentially affect receptor signalling, internalisation or cleavage and, therefore, that an appreciation of their binding sites is important. Hence, we decided to attempt to determine the crystal structures of TREM2 bound to scFv-2, scFv-3 and scFv-4 antibodies whose binding kinetics suggested strong interactions. Crystallisation screens were established using the TREM2 ectodomain (His19-Asp131), which has been structurally characterised previously (Kober *et al*., 2016; Sudom *et al*., 2018), as well as using the TREM2 ectodomain and stalk (His19-Ser174), which is relevant to shedding by ADAM proteases at H157, but susceptible to degradation *in vitro* due to the inherent flexibility of the C-terminal stalk region (Sudom *et al*., 2018). Both TREM2 proteins were produced in Expi293F™ human cells in the presence of kifunensine and deglycosylated using Endo H treatment.

Diffracting crystals were first obtained for the complexes of the smaller TREM2 ectodomain with both scFv-2 and scFv-4, but not scFv-3, enabling their structure determination at 3.36 and 3.07 Å resolution, respectively (see **Table 1** for data collection and refinement statistics). Both crystal lattices were characterised by a high solvent content (**Figure S2**), and despite screening over 180 crystals, the resolution of the scFv-2 complex could not be improved. Nevertheless, this level of resolution was sufficient to define the scFv-2/TREM2 interface. A higher resolution structure of the scFv-4 complex was, however, subsequently obtained using the longer TREM2 construct comprising both the ectodomain and part of the stalk region and refined to 2.26 Å resolution. Generally, the scFv-2/TREM2 complex contained fewer crystal contacts than scFv-4 leading to a more spacious lattice and possibly lower resolution (**Figure S2)**.

**Table 1.** Data collection and refinement statistics (molecular replacement)

|  | 6Y6C (scFv-4) | 6YMQ (scFv-4) | 6YYE (scFv-2) |
| --- | --- | --- | --- |
| **Data collection** |  |  |  |
| Space group | P 41 2 2 | P 21 21 2 | I 2 2 2 |
| Cell dimensions |  |  |  |
| *a*, *b*, *c* (Å) | 111.66 111.66 232.20 | 167.51 180.77 125.51 | 113.94 126.20 225.01 |
| $\alpha$, $\beta$, $\gamma$ (°) | 90 90 90 | 90 90 90 | 90 90 90 |
| Resolution (Å) | 100.63 - 2.26 (2.341 - 2.26) | 75.99 - 3.07 (3.18 - 3.07) | 110.07 - 3.36 (3.36-3.57) |
| *R*<sub>merge</sub> | 0.2129 (2.041) | 0.2811 (1.86) | 0.266 (1.55) |
| *I* / σ*I* | 9.96 (1.25) | 4.2 (1.0) | 7.4 (1.7) |
| Completeness (%) | 99.80 (99.34) | 99.83 (99.90) | 92 (68.9) |
| Redundancy | 21.4 (14.3) | 6.4 (6.6) | 13.4 (12.7) |
| Wilson B-factor | 39.44 | 69.70 | 68.7 |
| **Refinement** |  |  |  |
| Resolution (Å) | 2.26 | 3.07 | 3.36 |
| No. reflections | 69383 (6776) | 71714 (7075) | 16272 (70) |
| *R*<sub>work</sub> / *R*<sub>free</sub> | 0.191/0.216 | 0.2730/0.2871 | 0.298/0.328 |
| No. atoms |  |  |  |
| Protein | 5368 | 15625 | 5038 |
| Ligand/ion | 28 | 92 | 14 |
| Water | 375 | 13 | - |
| *B*-factors |  |  |  |
| Protein | 42.31 | 88.15 | 91.39 |
| Ligand/ion | 67.13 | 90.45 | 91.5 |
| Water | 47.65 | 58.05 | - |
| R.m.s. deviations |  |  |  |
| Bond lengths (Å) | 0.002 | 0.001 | 0.003 |
| Bond angles (°) | 0.51 | 0.40 | 0.56 |
| Solvent (%) | 73.85 | 69% | 77.18 |
| Average B-factor (Å<sup>2</sup>) | 42.0 | 88.14 | 91.0 |
| Anisotropy | 0.484 | 0.641 | 0.041 |

### ScFv-2 and scFv-4 bind to partially overlapping epitopes on TREM2 distal to the putative ligand-binding site

Here, we focus our subsequent work on the higher resolution structure of the scFv-4 complex with the TREM2 ectodomain and stalk (**Figure 3a**) and the scFv-2 complex with the TREM2 ectodomain alone (**Figure 3b**). In both structures, the globular ectodomain of TREM2 displayed a β-sandwich fold that consisted of 9 β-strands, including strands A-G common to all Ig-like domains and a C’-C’’ insertion typical of the V-set Ig domain (**Figure 3c**). The overall structure does not differ significantly to the previously solved structures of wild-type TREM2, PDBs 5ELI and 5UD7, except for some variability in the flexible loops **(Figure S3a).** The putative ligand-binding site on TREM2 is proposed to be located at one end of the β-sandwich, which presents the three complementarity-determining regions (CDRs1-3) and includes the AD-risk allele site Arg47 as part of CDR1 (Kober *et al*., 2016; Sudom *et al*., 2018). Of note, both scFv-2 and scFv-4 were found to bind to the opposite end of TREM2 (**Figure 3a-c**). Here, the β-sandwich of TREM2 splayed between β-strands C and F to provide an extended surface for the scFvs to bind that included parts of β-strands A, F, G and loop C-C’ (**Figure 3c**). However, the superposition of the two TREM2-scFv complexes revealed that the binding epitopes on TREM2 were only partially similar (**Figure 3c**). For instance, scFv-4 formed interactions with the C-C’ loop in TREM2 that were largely absent in the scFv-2 complex.

**Figure 3.**
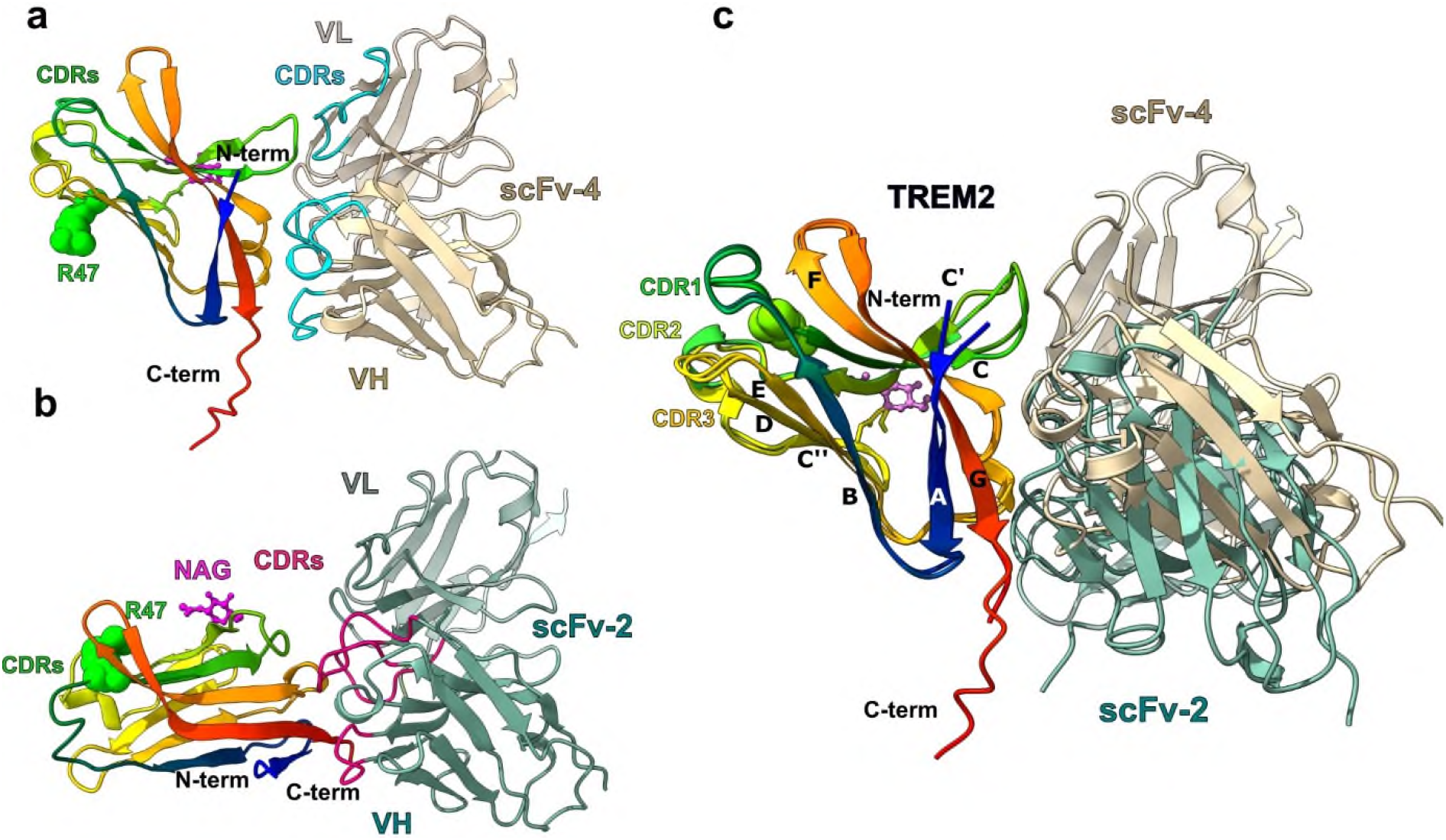
ScFv-2 and scFv-4 bind to partially overlapping epitopes on TREM2 distal to the putative ligand-binding site. (**a**) Cartoon representation of TREM2 bound to scFv-4. TREM2 β-sheets are labelled and coloured in rainbow. ScFv-4 is coloured in tan with VH domain positioned on the same plane as VL in lighter tan. CDR loops involved in TREM2 binding are coloured in cyan. R47 in CDR2 is coloured green in space fill, N-acetyl glucosamine remaining after de-glycosylation is coloured in pink and shown as ball and stick. (**b**) The general topology of TREM2 and scFv-2 complex. Panels (a) and (b) are fixed on the antibodies resulting in rotation of the TREM2 structures. VH domain of scFv-2 is coloured in green and VL domain is coloured in light green. CDR loops involved in TREM2 binding are coloured in magenta. (**c**) Overlay of both structures fixed on TREM2. The antibodies bind a similar epitope of TREM2 by engaging different CDR loops. The TREM2 epitope consists of strand βA (T17-A28), loop C-C’ (R52-P59) and residues in βF (P102-L107) and βG (L125-D131). ScFv-4 interacts extensively with the TREM2 C-C’ loop whereas scFv-2 interacts only with a fraction of the loop. A small change in the length of a short α-helix in the TREM2 CDR-2s is evident shown in green for scFv-4 and yellow for scFv-2 bound structures.

### Interactions of scFv-4 with the TREM2 ectodomain

Overall, the variable heavy (VH) and light domains (VL) of scFv-4 displayed canonical immunoglobulin folds tethered by an engineered glycine-serine rich linker that was disordered in the refined structure. The binding of scFv-4 to TREM2 was centred on the CDR3 loop of the scFv-4 VH domain (herein CDR-H3), flanked by interactions from CDRs-H1 and -H2 and by the VL domain CDR-L2. (**Figure 4a-b**).

**Figure 4.**
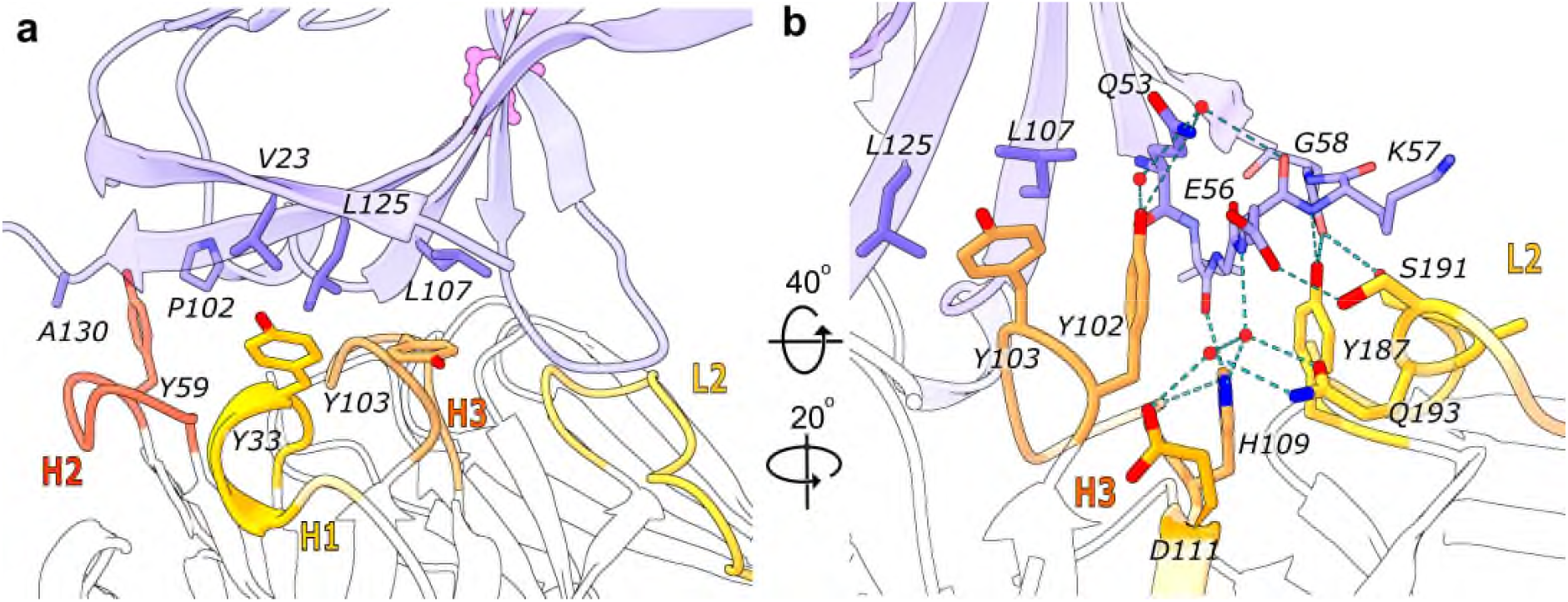
Interactions of scFv-4 with the TREM2 ectodomain. (**a**) Tyrosine residues of CDR-H1, H2 and H3 intercalating between TREM2 β-sheets rich in hydrophobic residues are highlighted as sticks. Interface residues are presented as blue (TREM2) or orange, red and yellow (scFv-4). (**b**) The same structure visualised by rotation as indicated. Residues involved in the hydrogen bond network between CDR-H3, CDR-L2 and TREM2 are presented as sticks. Only hydrogen bonds generated by scFv-4 side chains are shown together with water molecules mediating hydrogen bonding when applicable. A large number of contacts are with the TREM2 C-C’ loop. Calculated hydrogen bonds are shorter than 3.5 Å with angle 180°±10°.

The resolution of the scFv-4 complex structure enabled a detailed analysis of the side chain interactions in the binding interface (**Figure 4a-b**). A striking feature of the CDR sequence selections in scFv-4 was the enrichment of tyrosine residues (**Figure 1**). Indeed, CDR-H3 contained a sequence of six consecutive tyrosines, interrupted only by Gly104 (numbering corresponding to PDB entry sequences), which was likely selected to avoid the steric clashes that would result from any other amino acid at this position. Tyrosine residues 33, 59, 102, 103, 106, 107 and 187 in scFv-4 contributed to extensive hydrophobic packing interactions with the non-polar TREM2 residues Val23, Pro102, Leu107, Leu125, Val128 and Ala130 **(Figure 4a-b).** Additional side chain and water-mediated hydrogen bond interactions were provided by scFv-4 loops CDR-H3 and CDR-L2 (**Figure 4b**). The most extensive hydrogen bond interactions occurred between the CDR-L2 with the C-C’ loop of TREM2. Here, the scFv-4 residues Tyr187, Ser191 and Ser194 formed hydrogen bonds with TREM2 residues Lys47, Gly58 and Glu56, respectively (**Figure 4b**). Tyr102 from CDR-H3 also inserted between TREM2 β-strands C and F to form a further hydrogen bond with TREM2 Gln53 (**Figure 4b**). No significant differences were observed in scFv-4 binding to the TREM2 ectodomain in the equivalent structure using the shorter TREM2 construct His19-Asp131 (**Figure S3b**).

While the primary scFv-4 interaction was mediated by the TREM2 ectodomain as described above, we identified crystal contacts between the TREM2 stalk and CDR-H1 from a neighbouring scFv-4 subunit. This additional packing allowed the stalk to be traced in the electron density maps up to Asp137 in the C-terminus and several hydrogen bond interactions were observed to support the stalk interaction (**Figure S4)**

### Interactions of scFv-2 with the TREM2 ectodomain

The binding of scFv-2 to TREM2 was mediated by four CDR loops varied in the scFv library, including the heavy domain CDRs-H1, H2 and H3 and light domain CDR-L3 (**Figure 5a-b**). Overall, these loops were enriched in tyrosine and serine residues that contributed to a mix of hydrophobic and hydrogen bond interactions. Notably, TREM Glu127 (βG strand) was within hydrogen bonding distance of three scFv-2 side chains, including Ser54, Ser56 and Ser59 from CDR-H2. CDR-H2 Ser59 was additionally within hydrogen bonding distance of TREM2 Gln25 (βA). The sole glycine in CDR-L3 (Gly231) also contributed a hydrogen bond from its carbonyl group to the side chain of TREM2 His103.

**Figure 5.**
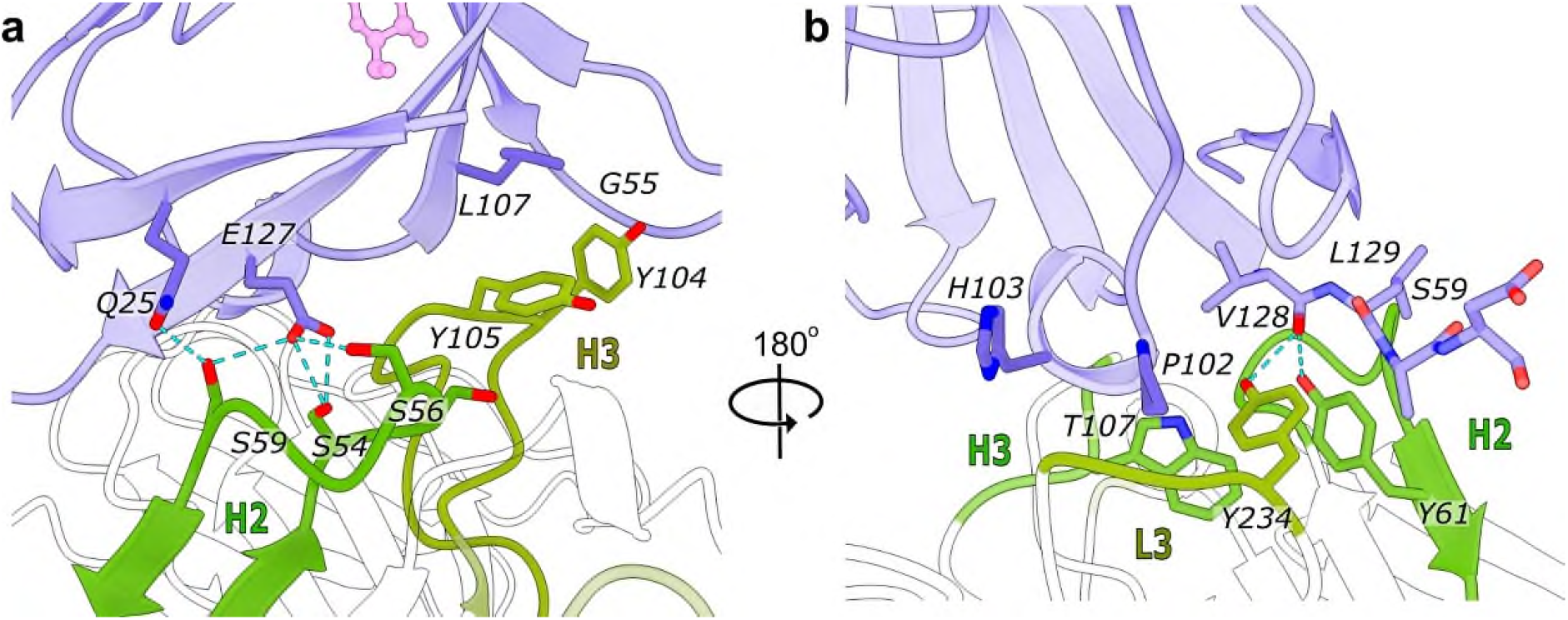
Interactions of scFv-2 with the TREM2 ectodomain. (**a**) Residues involved in hydrogen bonding and hydrophobic packing are represented as sticks and shown in blue (TREM2) or green (scFv-2). The aliphatic side chains of tyrosines 104 and 105 in CDR-H3 pack against the backbone of the C-C’ loop (Gly55) and Leu107 of TREM2. The serine rich CDR-H2 mediates multiple hydrogen bonds with the side chains of TREM2 Glu127 and Gln25. (**b**) 180°C rotated view of the interface between TREM2 and scFv-2. Hydrogen bonds are observed between the hydroxyl groups of Tyr234 (L3) and Tyr61 (H2) and the carbonyl of TREM2 Val128. Further hydrophobic packing is mediated by Trp107 (H3) and other side chains from CDR-H2 and CDR-L3.

Together the scFv-2 and scFv-4 complex structures show the potential value of these antibody fragments as crystallisation chaperones. More importantly, however, they define the epitopes of these scFvs, allowing for the interpretation of functional properties in the context of the binding site on the surface of TREM2, which has been poorly described for other anti-TREM2 antibodies.

### ScFv-3 and scFv-4 reduce shedding of sTREM2 from HEK293 cells overexpressing TREM2 and DAP12

The H157Y polymorphism, which leads to accelerated cleavage of TREM2 and increases AD risk, suggests that regulation of shedding of sTREM2 might be important in human biology (Jiang *et al*. 2016; Thornton *et al*. 2017). Based on the potential interactions of scFv-4 with the stalk domain as indicated by the crystal contacts, and the potential for TREM2 internalisation induced by scFv dimers, we decided to determine the effect of the scFv antibodies on sTREM2 shedding. We devised a sandwich ELISA, capturing sTREM2 from cell culture supernatants, and confirmed that scFvs were not masking the sTREM2 epitopes required by the ELISA capture and detection antibodies **(Figure S5**). Recently another metalloprotease, meprin β, was shown to cleave TREM2, with the main cleavage site between Arg136 and Asp137 (Berner *et al*., 2020). We confirmed that both the ectodomain as well as a sequence of stalk C-terminal to Ala138 were required for detection in our ELISA protocol. Thus, the detected sTREM2 product is unlikely to be resulting from meprin β cleavage **(Figure S5**). Batimastat, a broad spectrum chemical inhibitor of matrix metalloproteinases, including ADAM proteases, was used as a positive control to inhibit shedding. HEK293 cells stably transfected with TREM2 and DAP12 were treated with each of the scFvs for 5 hours to reach the appropriate threshold for sTREM2 detection in the supernatant. ScFv-3 and scFv-4 (10 μg/mL) significantly reduced shedding of sTREM2 by 22% and 38%, respectively (**Figure 6a**).

**Figure 6.**
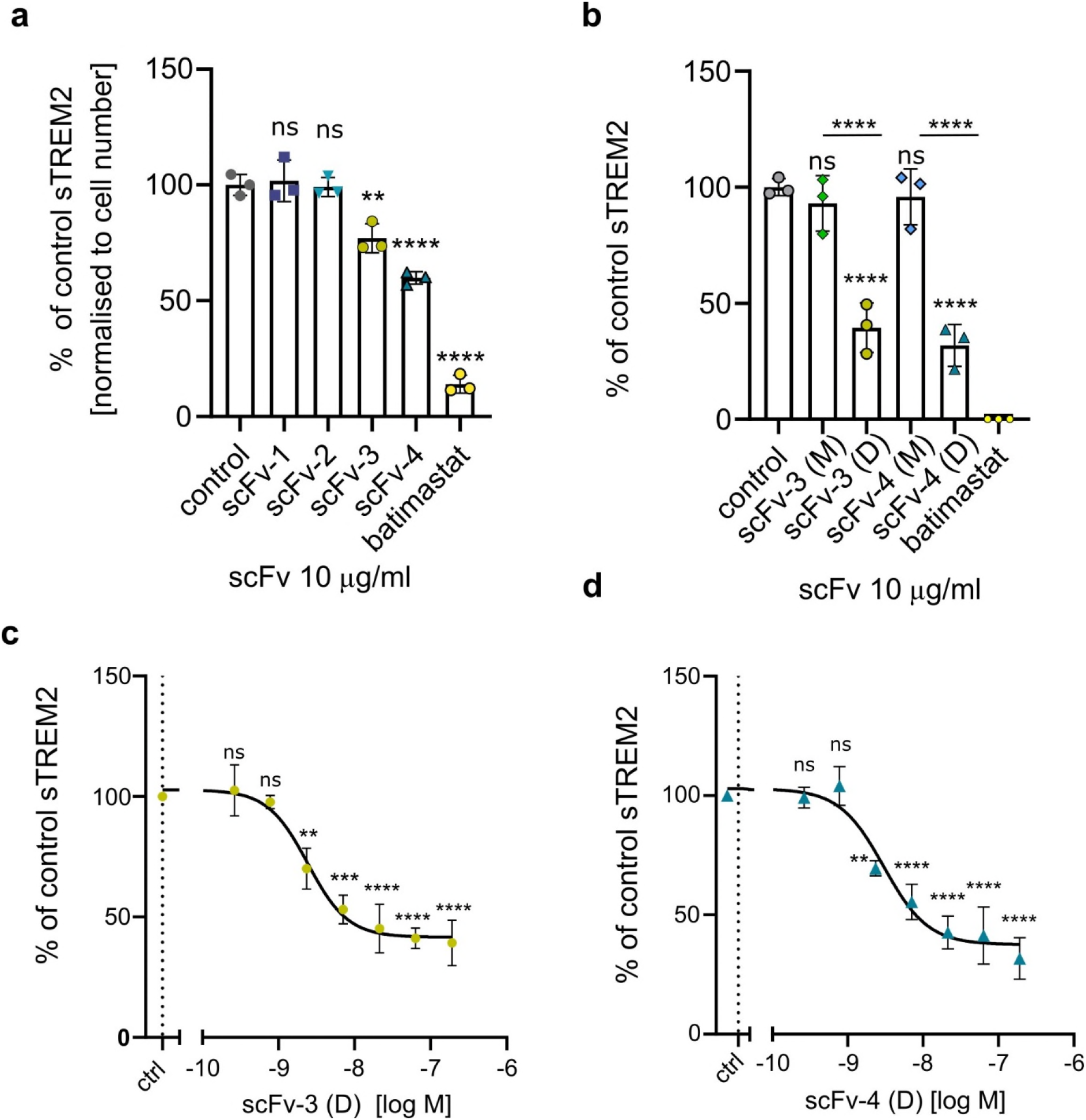
ScFv-3 and scFv-4 and their oligomers reduce TREM2 shedding in HEK293 cells overexpressing TREM2 and DAP12. (**a**) HEK293 cells overexpressing TREM2 and DAP12 were treated with scFvs at 10 μg/mL for 5 hours or with 20 μM batimastat or buffer in control samples. Data were normalised to cell number by nuclei counting and represent the mean ± SEM (n=3). One-way ANOVA, Dunnett’s post hoc test comparison to buffer only control was carried out with significant P values scFv-3 vs control=0.0011, scFv-4 vs control <0.0001, batimastat vs control < 0.0001. (**b**) Results of shedding assay described above for separated scFv monomers (M) or dimers (D). One-way ANOVA, Dunnett’s post hoc test comparison with significant P values scFv-3 (D) vs control < 0.0001, scFv-4 (D) vs control < 0.0001, monomer versus dimers < 0.0001, sTREM2 in batimastat samples were below detection threshold. (**c**) Assay of scFv-3 dimer showing scFv-concentration-dependent reduction in shedding. Data represent n=3 independent experiments, One-way ANOVA, Dunnett’s post hoc comparison to buffer control, test P < 0.0001, EC50=2.3 nM (**d)** scFv-4 dimer concentration-dependent reduction in shedding. Data represent n=3 independent experiments, One-way ANOVA, Dunnett’s post hoc comparison to buffer control, test P < 0.0001, EC50=3 nM.

Oligomerization of scFvs has been observed and can be exploited to enhance the properties of selected molecules (Kortt *et al*., 2001). We decided to investigate the effect on shedding of the apparent monomer and dimer species present in our scFv-3 and scFv-4 samples. After these species were acutely separated by SEC and processed rapidly to minimise re-equilibration between low and high molecular weight (MW) species, it was found that the monomer species for both scFv-3 or scFv-4 had minimal or no activity **(Figure 6b)**, whereas the dimers decreased production of sTREM2 by more than 50% **(Figure 6b).** Concentration-response experiments for the scFv-3 and scFv-4 dimers showed that maximum effect plateaued between 60-70% with calculated EC50 values of 2.3 nM and 3 nM, respectively **(Figures 6c-d).** Of note, scFv-2, which showed a high binding affinity for TREM2, caused no changes in sTREM2 release despite having a similar binding epitope to scFv-4 (**Figure 6a**). This antibody fragment purified mostly as a monomer, further highlighting the importance of scFv oligomerization for reduction of shedding. None of the scFvs impacted cell viability, as measured by counting cell nuclei using Hoechst staining and fluorescence-based viability (data not shown).

Finally, in order to demonstrate selective binding to TREM2, we conjugated the dimeric fractions of scFv-3 and scFv-4 to Alexa fluor 647 and repeated the HEK293 treatment protocol with subsequent imaging of the fluorescent scFvs. Labelling of the cells was dependent upon the heterologous expression of TREM2/DAP12. The majority of Alexa signal was present intracellularly and was resistant to acidic wash, consistent with TREM2 mediated internalisation under the conditions tested. ScFv-4 appeared to show superior specificity compared to scFv-3 based on the staining of non-transfected HEK293 cells **(Figure 7).**

**Figure 7.**
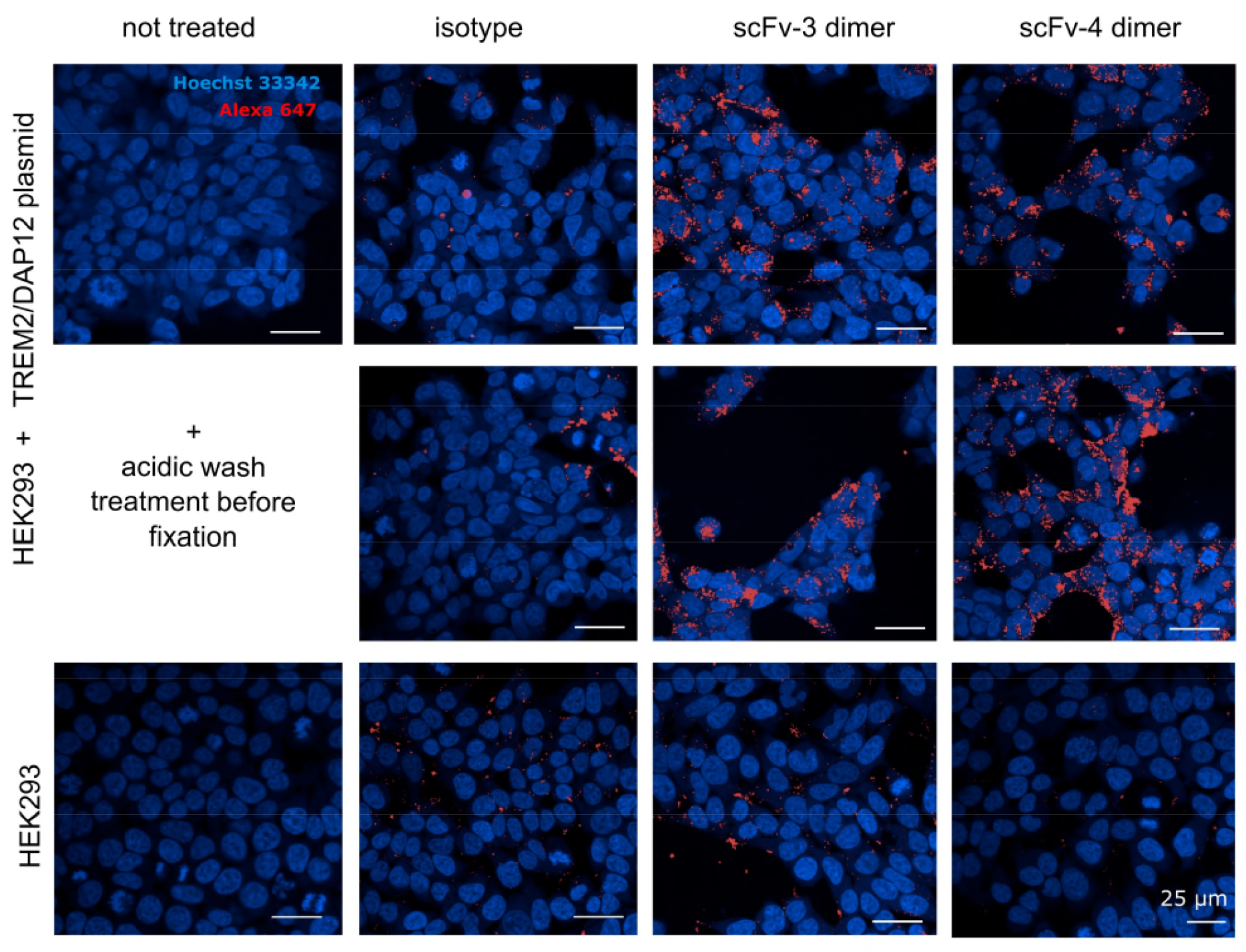
ScFv-3 and 4 engage TREM2 in HEK cells overexpressing TREM2/DAP12 and produce intracellular signal after 5-hour treatment. HEK293 cells transfected with TREM2/DAP12 or WT HEK293 were incubated with dimers of scFv-3 and scFv-4 or control scFvs against different target pre-conjugated to Alexa fluor 647 or buffer for 5 hours. Cells were washed with medium or DMEM pH 2 medium twice before fixation and staining. Nuclei were visualised by Hoechst staining (blue) and scFvs by Alexa fluor 647 (red) by OperaPhenix confocal lens with 40x magnification, n=1 with 3 technical repeats. Scale bar denotes 25 μm.

## Discussion

In this paper, we have presented our work on the structural characterisation of scFvs raised against human TREM2 by phage display. We screened a library of scFv antibody fragments against fully folded and glycosylated TREM2 ectodomain which resulted in the production of three successful candidates with good binding affinities. Despite challenging purification procedures, we were successful in producing recombinant antibody fragments and demonstrated them to be suitable for X-ray crystallography. We have solved the structures of two scFvs bound to TREM2 and provide the first crystallographic structures of antibody interactions with TREM2. These are also the first antibody fragments reported to unambiguously bind TREM2 19-131 globular ectodomain. Additionally, we have shown that higher molecular weight oligomeric species of scFvs are able to display enhanced functional activity, suggesting parallels with the activity of bivalent mAbs.

Since TREM2 is a potential therapeutic target for AD, the pharmaceutical industry has initiated several projects aimed at developing TREM2 antibody therapeutics, with at least two candidates having commenced clinical trials. Data have been reported for two, AL002 and 4D9, which allows for an interesting comparison with scFv-4 (Schlepckow *et al*., 2020; Wang *et al*., 2020). Other functionally active antibodies have either not been widely published (WO2016023019A2) or the epitope has not been defined and, therefore, are not included in the discussion below (Cheng *et al*., 2018). X-ray crystallographic data, which accurately define the antibody/TREM2 interfaces, are not publicly available for any of these reagents. AL002 is an agonist antibody with a poorly defined epitope most likely in the stalk domain between residues 112-174 (Wang *et al*., 2020), while 4D9 has an epitope defined by peptide mapping to residues Asp137-Gly145 (Schlepckow *et al*., 2020). As such, they target TREM2 epitopes N-terminal of the α-secretase cleavage site at H157, whereas our X-ray crystallography data shows that scFv-4 binds the immunoglobulin domain of TREM2. The binding sites do show some cross-species variation explaining the observation that 4D9 is specific for mouse TREM2, whereas AL002 has a high affinity for human TREM2.

As both AL002 and 4D9 are stalk binding antibodies it is possible that they might affect cleavage of TREM2 by virtue of sterically hindering access by ADAMs. Consistent with that hypothesis, both full-length antibodies (4D9 and AL002) have been shown to reduce shedding of sTREM2. This interesting property has been confirmed for the first time in human with AL002 (Wang *et al*. 2020). This function also appears to be dependent upon the use of a bivalent antibody, as the 4D9 Fab fails to inhibit shedding of sTREM2 by cells. These observations are in agreement with our data, where higher MW oligomers of scFv-3 and scFv-4 inhibit up to 70% of sTREM2 production.

At present, the mechanism of action for the inhibition of shedding by all of these antibody reagents is unclear. It might occur either by inhibition of cleavage of TREM2 at the cell surface or by promoting internalisation of full-length TREM2, thereby removing substrate availability. Schlepckow *et al*. propose that 4D9 exerts some of its effects by stabilising TREM2 at the plasma membrane, explaining the potentiation of signalling in response to liposomes by 4D9. However, it is also possible that reduction of sTREM2 production by bivalent antibodies or oligomeric scFvs is due to cross-linking of receptor and internalisation, thereby reducing cell surface substrate for shedding. It is noteworthy that a bivalent antibody is necessary for 4D9 functional effects and to date we have not observed any functional responses to monomeric scFv-4, only observing activity in preparations containing appreciable dimeric species. Furthermore, Alexa Fluor labelled scFv-4 and scFv-3 label HEK293 cells in a TREM2 expression dependent manner, leading to intracellular immunofluorescence, although we have not yet demonstrated that this is endocytosis dependent. Interestingly, the scFv-2 antibody fragment has the lowest propensity to form oligomeric species and did not show any inhibitory activity in the sTREM2 shedding assay. Our data is therefore consistent with the effect of scFv-3 and −4 upon shedding of sTREM2 being due to promoting the internalisation of TREM2.

Both AL002 and 4D9 show agonist properties *in vitro*. AL002 elicits TREM2 mediated secondary intracellular signalling, demonstrated using an NFAT reporter cell line and phosphorylation of a protein of equal molecular weight to DAP12 in bone-derived macrophages (Wang *et al*. 2020). Similarly, 4D9 shows agonism, but only when a full-length antibody was used to challenge cells; Syk phosphorylation did not occur following exposure of cells to 4D9 Fab fragment. The requirement for a bivalent antibody for the function of AL002 has not been reported, but these observations with 4D9 led the authors to hypothesise that the functional effects of 4D9 are dependent upon a bivalent antibody and cross-linking of TREM2 molecules (Schlepckow *et al*. 2020). Taken together, the literature and our own data highlight the importance of multivalency for the functional activities of anti-TREM2 antibodies. On account of the need to prepare bivalent mAb and Fab antibodies, a full investigation of the agonist and antagonist properties of the scFvs obtained in our studies is the subject of a separate piece of work.

While we have presented evidence that scFv capable of dimerization reduce sTREM2 shedding from the HEK293 cells, the functional impact of monomeric scFvs is unclear due to the dynamic equilibrium between monomeric and dimeric forms. However, we have observed a decrease in activity upon storage of a purified fraction of dimers and also a lack of activity of freshly prepared and rapidly utilised monomers, suggesting that monomer represents an inactive conformation (data not shown). Similarly to the mAbs discussed above, multivalent scFvs will have the potential to cross-link TREM2 ectodomains into higher-order structures, thereby creating significant steric hindrance to proteolytic complexes, altering their characteristics as substrates, or triggering endocytosis.

By using phage display to raise recombinant scFvs we have assured that the source of the molecules is renewable and can be easily shared. During our studies, we have encountered difficulties caused by the use of commercially available antibodies with unknown TREM2 epitopes, especially when used in ELISA, pull-down studies or immunocytochemistry. The presented scFvs bind away from the proposed ligand recognition site, qualifying them for development into additional tools for the study of TREM2-ligand interactions and receptor processing.

Our data represent the first crystallographic characterisation of binding interactions between functional antibody fragments and TREM2 and we hope that these additional tools will help elucidate the complex pathophysiology of TREM2 in Alzheimer’s disease.

## Acknowledgements

The authors would like to thank Diamond Light Source for beamtime (proposal mx15433), as well as the staff of beamLine I24 for assistance with crystal testing and data collection. We would like to acknowledge Prof. Peter St George-Hyslop for the gift of TREM2-DAP12 expressing HEK cells and valuable advice, Andy Takle, James Staddon, Karel Otero-Gutierrez and Suresh Poda for advice during the project. The SGC is a registered charity (number 1097737) that receives funds from AbbVie, Bayer Pharma AG, Boehringer Ingelheim, Canada Foundation for Innovation, Eshelman Institute for Innovation, Genome Canada, Innovative Medicines Initiative (EU/EFPIA) [ULTRA-DD grant no. 115766], Janssen, Merck KGaA Darmstadt Germany, MSD, Novartis Pharma AG, Ontario Ministry of Economic Development and Innovation, Pfizer, São Paulo Research Foundation-FAPESP, Takeda, and Wellcome Trust [106169/ZZ14/Z]. AS was supported by a DPhil studentship sponsored by Eisai. JBD, EM, TBS and EDiD were supported by a grant from Alzheimer’s Research UK (ARUK-2015DDI-OX).

## Author contributions

Conceptualisation AS, JBD, NABB, ANB; provision of resources YC, JY, DQ, SM, SG, PJA, JW; investigation AS, JY, DQ, CP, EW, supervision YC, TBS, SG, HP, PJA, EDiD, EM, JBD, NABB, ANB; manuscript preparation AS, EDiD, JBD, NABB, ANB; funding acquisition JW, JBD. All authors reviewed the submitted manuscript.

## Competing interests

YC, SJN, PJA and JW are employees of Eisai Ltd., and Eisai Inc.

## Resource availability

### Lead Contact

JBD

### Materials Availability

Plasmids generated in this study will be deposited to Addgene and made available on request.

### Data and Code Availability

The coordinates and structure factors for the crystal structures reported in this article have been deposited in the PDB with accession codes 6YYE (TREM2-scFv-2) and 6Y6C (TREM2-scFv-4), 6YMQ (TREM2-scFv-4).

## STAR Methods

### Cloning of TREM2 and scFvs

All plasmids were generated in MACH1 cells or DH5α (Thermofisher Scientific) and bacmid DNA for insect cell expression and baculovirus generation were generated in DH10Bac (Thermofisher Scientific).

#### Cloning of TREM2 for ELISA validation

DNA encoding human TREM2 (amino acids 19-131; gi|9507203; MGC) used for antigen generation and TREM2 (a.a. 19-138) for ELISA validation were cloned into the baculovirus secretion plasmid pFB-Sec-Bio5 which provides a C-terminal avi-tag sequence for biotinylation. The stalk-containing TREM2 (a.a. 19-174) used for ELISA validation was cloned into baculovirus secretion plasmid pFB-Sec-NH which provides a N-terminal hexahistidine tag.

#### Cloning of TREM2 for SPR

A TREM2 construct containing the native signal peptide (a.a. 1-174) was cloned into the pTT5 vector, followed by an avi tag and TEV-cleavable hexahistidine tag (GLNDIFEAQKIEWHEGSENLYFQSHHHHHH) at its C-terminus.

#### Cloning of TREM2 for X-ray crystallography

For structural studies, human TREM2 (a.a. 19–131) was cloned into the mammalian expression vector pTT5, following an N-terminal signal peptide (MGILPSPGMPALLSLVSLLSVLLMGCVAETG) with hexahistidine tag preceeded by additional linker (a.a. GTK) at its C-terminus.

The longer construct TREM2 (a.a. 19-174) containing the stalk was inserted by ligation independent cloning (LIC) (Strain-Damerell *et al*., 2014) into the vector pHTBV1.1-SecNH-Bio, which provides an N-terminal Gp64 signal peptide and TEV-cleavable hexahistidine tag as well as a C-terminal avi tag.

#### Cloning of scFvs for baculovirus expression

ScFv antibody fragments identified by phage display were cloned for baculovirus expression with scaffold specific primers into the vector pFB-Sec-NH.

### Expression and purification of TREM2 and scFvs in baculovirus expression system

#### Bacmid generation and transfection

TREM2 viruses for baculovirus expression were generated by transfection with bacmid DNA:JetPrime (Polyplus) complexes. Sf9 cells at a density of 0.2×10^6^ in adherent culture were transfected for 4 hours (h) at 27°C followed by medium exchange to SF900 II-SFM containing 0.1% penicillin and streptomycin (Thermofisher Scientific) with 2% FBS (Thermofisher Scientific). P0 viruses were harvested after 7 days at 27°C and 120 μL was used to infect 3 mL of Sf9 cells for P1 virus amplification (72 h and 450 rpm shaking and throw 6 mm). Following another round of viral amplification, 0.75 L of Sf9 in SF900 II-SFM media were infected with virus P2 at 7 mL/L at density 2×10^6^ in glass bottles (27°C, 170 rpm for 72 h, throw 25 mm).

#### Purification of TREM2 for antigen generation and ELISA validation

The clarified supernatants were loaded onto 5 mL Ni-NTA (Qiagen)/per 1L at 10 mL/min. Resin was washed with 20 column volumes (CV) of 50 mM HEPES pH 7.4 and 300 mM NaCl, 5% glycerol, 5 mM imidazole followed by 10 CV of 10 mM imidazole and 600 mM NaCl containing buffer. Proteins were eluted with buffer containing 500 mM imidazole. Proteins were purified further by size exclusion chromatography (Superdex 200,16/60) *in vitro* biotinylated using recombinant BirA enzyme (enzyme to substrate ratio of 1:50, w/w). The reaction was performed in a buffer containing 10 mM Tris-HCl, 50 mM NaCl and 10 mM MgOAc, pH 8.0 for 14 h, at 4°C in the presence of 0.1 mM biotin and 10 mM ATP. Subsequently, the buffer was exchanged by dialysis to 20 mM HEPES, 300 mM NaCl, 5% glycerol and 0.5 mM TCEP, pH 7.5

TREM2 (His19-Ala138) and (His19-Ser174) for ELISA validation were purified in the same manner but were stored at −80°C after size exclusion.

#### Expression and purification of scFvs

Viruses for scFvs were generated as above. IMAC was performed as for TREM2, except for a change to pH 6.8 which decreased precipitation. Eluted proteins were cleaved with TEV protease overnight (O/N) at 4°C followed by loading onto a 1 mL Protein A FF (GE healthcare) column at 3 mL/min. Resin was washed with 20 CV of base buffer (BB: 25 mM HEPES pH 7.5, 150 mM NaCl) followed by a wash with 10 CV of BB containing 0.05% Triton-X. Proteins were eluted with 0.1 M acetic acid pH 3 at 0.5 mL/min and immediately buffer exchanged on PD10 columns (scFv-1: 120mM Citrate pH 7 with 50 mM L-arginine,50 mM L-glutamic acid, scFv-2: 100 mM HEPES pH 7.2, 25 mM NaCl, 100 mM L-arg, 100 mM L-glu, scFv-3: 100 mM Tricine pH 8, scFv-4: 100 mM Tricine pH 8) and frozen until further use. Later, scFv-3 and scFv-4 oligomers were separated by double SEC on Superdex 200 16/60 in 20mM Tris-HCl pH 7.5, 300mM NaCl and 5% glycerol. Separated oligomers were pooled and buffer exchanged to PBS using vivaspin 2 10 MWCO concentrators (GE healthcare). Proteins were clarified by centrifugation at 30,000 x *g* and immediately frozen in liquid nitrogen.

### Expression and purification of TREM2 in mammalian cells

#### Transient transfection of TREM2 plasmids

For crystallisation studies of TREM2 (a.a. 1-131), Expi293F™ cells (Thermofisher Scientific) in Expi293™ expression medium (Thermofisher Scientific) at 2.5×10^6^ were transfected with a mixture of 1 mg of pTT5:TREM2 (19-131) DNA to 6 mg L-PEI 25 K (Polysciences) per 1 L of culture. 50 mL of DNA and 50 mL of L-PEI in Opti-MEM (Thermofisher Scientific) was mixed and incubated for 15 min before addition to the culture. 1 mg/L kifunensine (Carbosynth) was added followed by 1 mM valproic acid (VPA), 20 h posttransfection. Cells were grown in vented flasks at 100 rpm (throw 25 mm), 37°C, 8% CO2 for 72 h.

#### Purification of TREM2 for crystallisation

Culture supernatants were collected and TREM2 (a.a 19-131) was purified by affinity chromatography (Ni-NTA, Qiagen) in buffers containing 20 mM Tris-HCl pH 8, 500 mM NaCl, 5% glycerol followed by size exclusion chromatography (Superdex 75 10/30GL, GE Healthcare). The purified protein was deglycosylated with EndoH (PE019, Novoprotein) in 50 mM sodium citrate pH 5.5 overnight at 4°C, at the ratio of 1 mg protein: 2000 U EndoH. EndoH was removed using a second Ni-NTA affinity column, and the purified protein was stored in buffer containing 50 mM Tris pH 8.5, 200 mM NaCl, and 5% glycerol at 10 mg/ml concentration.

The longer TREM2 (a.a. 19-174) for crystallisation was expressed and purified similarly but transfection was performed with DNA to L-PEI 1:3 ratio and cells were grown in 2 L roller bottles (Greiner Bio-one). Affinity chromatography was performed in base buffer: 20 mM HEPES 7.4, 300 mM NaCl, 5% glycerol and proteins were immediately deglycosylated in solution adjusted with sodium citrate (final buffer pH 6). Size exclusion was performed in 20 mM HEPES 7.4, 150 mM NaCl, 5% glycerol on Superdex 200 16/60 (GE Healthcare).

#### Purification of TREM2 for SPR

Fully glycosylated biotinylated pTT5-TREM2 (19-174) protein for SPR studies was purified as described for TREM2 (a.a. 19-131) without the deglycosylation step. The Ni-NTA purified protein was biotinylated overnight at 4°C in the presence of 0.5 mM biotin, BirA (1:50), 1 mM ATP, and 7.5 mM MgCl_2_. TREM2 was further purified by a second Ni-NTA affinity column and stored in buffer containing 50 mM Tris pH 8.5, 200 mM NaCl, and 5% glycerol.

### Antibody selection and validation

Antibody generation through phage display selection was performed as described previously (Preger *et al*., 2020), using the *in vitro* biotinylated TREM2A (19-131) as antigen and a human synthetic single-chain fragment variable (scFv) library. Briefly, four rounds of selections were performed using streptavidin beads (Dynabeads M-280 streptavidin, Invitrogen) to immobilise the biotinylated TREM2 antigen. In the first two rounds, phages were incubated with immobilized antigen after a pre-selection on naked beads. In round 3 and 4, phage and antigen were incubated in solution before capturing on beads. The antigen-phage incubation time was decreased from 3 h in the first round to 1.5 h in rounds 2, 3, and 4. Furthermore, the selection pressure was augmented by increasing the number of washing steps and decreasing the amount of antigen added (200, 100, 50 and 10 pmol, respectively) between the different rounds. The enriched phages were recovered using trypsin digestion and amplified overnight by infection of *E. coli* XL1-blue. Amplification in round 1 was done using agar plates and in rounds 2, 3 and 4 in solution. Amplified phages were precipitated using PEG/NaCl and used for the next round of selection. Phagemid DNA from round 3 and 4 were purified and the scFv genes were transferred to an expression vector as described previously (Preger *et al*., 2020).

Following selections, cloning and transformation, a total of 188 colonies were picked from round 3 and 4 and analysed further. Primary validation was performed as described including ELISA, HTRF, Luminex, and SPR (Preger *et al*., 2020). Based on these results, four scFv candidates, scFv-1, scFv-2, scFv-3, scFv-4, were selected to be produced in large-scale and tested in additional binding kinetics and co-crystallisation trials.

### ScFv binding kinetics

Approximate binding kinetics of Protein A purified ScFv-2, scFv-3 and scFv-4 were determined by SPR performed on a Biacore 8K instrument. ScFv-1 was previously determined to be binding TREM2 in the μM range and thus was not analysed in detail. Biotinylated TREM2 (19-174) produced in mammalian cells was immobilised on a streptavidin-coated chip supplied in the Biotin CAPture kit (28-9202-34, GE Healthcare) at 0.5 μg/mL concentration. ScFvs were diluted in 10 mM HEPES, 150 mM NaCl, 0.05% P-20 and injected at a flow rate of 30 μL/min for 60 seconds followed by dissociation for 120 seconds. The binding response was calculated after subtracting signal coming from a blank flow cell and the buffer. Langmuirian 1:1 model was used to calculate the binding kinetics. For scFv-3 and scFv-4 avidity contributed to obtained values. Additionally, scFv-4 showed some non-specific interaction with the reference sensor, thus so calculated values are used as approximation only.

### Crystallisation

#### scFv-4 PDB: 6YMQ

After Protein A purification, scFv-4 in 100 mM Tricine, pH 8 and TREM2 (a.a. 1-131) were mixed at molar ratio 1:1.1 and buffer exchanged into 20 mM Tricine pH 8, 200mM NaCl, 200 mM L-arginine, 200mM L-glutamic acid. The complex was concentrated to 14 mg/mL and crystals were grown by sitting drop vapour diffusion in solution containing 39% MPD, 0.2 M ammonium acetate, 0.1 M citrate pH 5.5 at 20°C. Crystals were mounted directly from the drop and vitrified in liquid nitrogen.

#### scFv-4 PDB: 6Y6C

Antibody scFv-4 was buffer exchanged into 25 mM HEPES, 300 mM NaCl and 5% glycerol and mixed with TREM2 (a.a. 1-174) at molar ratio 1:1.1. The complex was concentrated to 12.5 mg/mL and crystal plates were set up after 32,000 x *g* spin for 20 min. Protein crystals appeared after 1 day using a reservoir solution containing 0.2 M potassium chloride, 35% pentaerythritol propoxylate 5/4, 0.1 M HEPES pH 7.5 at 20°C. Crystals were cryo-protected after addition of reservoir solution containing 20% ethylene glycol and vitrified in liquid nitrogen.

#### scFv-2 PDB: 6YYE

Protein A purified antibody scFv-2 in 100 mM citrate buffer and 250mM L-arg and 50mM L-glu was mixed with TREM2 (a.a 19-131) at a molar ratio 1.3:1 and buffer exchanged on PD10 column to 60 mM Tricine 8, 200 mM NaCl containing 200 mM L-arg/L-glu solution. Complex was concentrated to 16 mg/mL and precipitation was removed by centrifugation. Crystals were grown in 15 mM nickel chloride, 0.1 M Tris pH 8.5, 2.2 M ammonium acetate at 20°C. Crystals were cryo-protected with 20 % ethylene glycol before vitrification.

### Data collection and structure determination

For scFv-4 structures (6YMQ, 6Y6C) data were collected on beamline I24 at the Diamond Light Source using the X-ray wavelength of 0.9688Å and processed using Xia2 package and DIALS (G. Winter 2010; 2018).

For the lower resolution scFv-2 dataset (6YYE), data were collected on beamline I24 at the Diamond Light Source using the X-ray wavelength 0.9763 Å. The initial model was built using PhenixRefine and coot modelling using Xia and DIALS dataset. Later the model was used to refine against AutoProc processed dataset with ellipsoidal truncation by Staraniso (Evans, 2006; Kabsch *et al*., 2010; Vonrhein *et al*., 2011, 2018). The model was finished by refinement in Buster v.2.10.3 (Bricogne *et al*., 2016) and final rounds of Phenix refine (Liebschner *et al*., 2019).

The first dataset of scFv-2 (6YYE) was phased using the structure of another scFv (PDB: 6G8R) generated from the same library and solved previously (Fairhead *et al*., 2019). TREM2 was phased using PDB 7UD5 (Sudom *et al*., 2018). The resulting scFv-2 co-structure (PDB 6YYE) was used for phasing of PDB 6YMQ. Phenix molecular replacement was used to phase the data and coot to build a model (Emsley *et al*., 2010; Liebschner *et al*., 2019). Models were improved by iterative cycles of Phenix Refine and coot rebuilding.

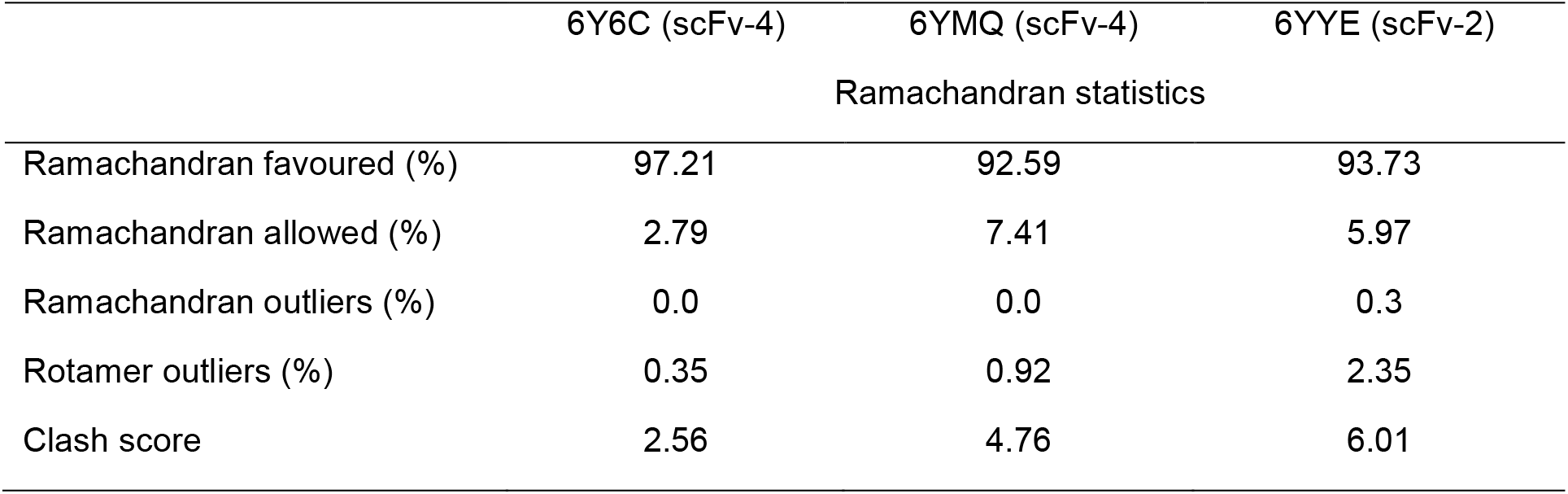

|  | 6Y6C (scFv-4) | 6YMQ (scFv-4) | 6YYE (scFv-2) |
| --- | --- | --- | --- |
| Ramachandran statistics |  |  |  |
| Ramachandran favoured (%) | 97.21 | 92.59 | 93.73 |
| Ramachandran allowed (%) | 2.79 | 7.41 | 5.97 |
| Ramachandran outliers (%) | 0.0 | 0.0 | 0.3 |
| Rotamer outliers (%) | 0.35 | 0.92 | 2.35 |
| Clash score | 2.56 | 4.76 | 6.01 |

Molecular graphics and analyses were performed with UCSF ChimeraX, developed by the Resource for Biocomputing, Visualization, and Informatics at the University of California, San Francisco, with support from National Institutes of Health R01-GM129325 and the Office of Cyber Infrastructure and Computational Biology, National Institute of Allergy and Infectious Diseases (Goddard *et al*., 2018). Crystal lattice images were generated using PyMOL (Schrödinger, LLC.).

### Mammalian cell culture for cell biology

The HEK293 TREM2-DAP12 cell line was a kind gift from Professor Peter St-George Hyslop and generated as described previously (Thornton *et al*., 2017). HEK293 wild type cell line was purchased from ATCC (CRL1573).

WT HEK293 and HEK293 cells stably expressing WT hTREM2 and hDAP12 were cultured in T-175 flasks (Greiner Bio-one) in DMEM (GIBCO) with 10% heat-inactivated foetal bovine serum (FBS) (Gibco) at 37°C, 5% CO2. The cell culture medium for the HEK293 TREM2-DAP12 cells was supplemented with 1 mg/mL puromycin hydrochloride (Cayman Chemical).

### sTREM2 ELISA

HEK293 transfected with TREM2-DAP12 were plated in 96-well plates (Corning Biocoat, 354640) at 30×10^3^ cells per well, grown for 24 h, washed with culture medium and treated with antibodies for 5 h. After antibody incubation, the medium was collected and transferred into a new plate, centrifuged at 1,000 x *g* followed by supernatant transfer into a fresh plate that was stored at −80°C.

TREM2 ELISA was performed by coating high bind 96-well plates (Sigma, M4436-040EA) with 8 μg/mL TREM2 antibody (Abcam, ab209814) O/N at 4°C followed by 1 h blocking using PBS with 1% BSA (Sigma). 100 μL of the defrosted cell supernatants were added at the appropriate dilution in DMEM + 1% BSA alongside a standard curve with TREM2 peptide (Sino Biologicals, 11084-H08H) and incubated for 2 h, at RT. 25 μL of the biotinylated detection antibody (R&D, BAF1828) at 1.5 μg/mL was added for an hour followed by the addition of HRP conjugated streptavidin diluted at 1:30,000 (ThermoFisher Scientific, 34028). TMB ELISA substrate (ThermoFisher Scientific) was added for 20 min followed by 2 M sulphuric acid (Immuno Chemistry technologies). Results were obtained by measuring optical density (OD) at 450 nm using a plate reader (SpectraMax M2). OD measurements in the linear range (0.8-1.9) were chosen for analysis and TREM2 concentration was interpolated from the TREM2 standard curve. To ensure that scFvs binding to TREM2 did not interfere with the assay antibodies, the standard peptide (a.a 1-174, SynoBiological) was mixed with each one of the scFvs in DMEM +1% BSA and results were compared to the standard curve only. Also, as another control, to assess TREM2 epitope recognition by the ELISA antibodies, TREM2 proteins (His19-Ala138 and His19-Ser174) were diluted to 2441 pg/mL in DMEM +1% BSA using five-point serial dilutions and analysed using the protocol described above. Additionally, the medium conditioned in cells for 5 h was spiked with scFvs and ELISA was performed in comparison to non-spiked sample (data not shown).

### Alexa fluor 647 cell staining and imaging

Purified scFv-3 and scFv-4 oligomers and control non-TREM2 scFvs were incubated with Protein A and Alexa Fluor 647 NHS ester (Thermofisher Scientific) at 1:10 molar ratio for 20 h, at 4°C and excess dye was removed by NAP5 columns (GE healthcare). HEK293 and HEK293 transfected with TREM2-DAP12 were prepared as for the ELISA and treated with scFv-4 for 5 h. Cells were fixed in 4% paraformaldehyde in PBS (Santa Cruz Biotechnology) for 15 min and kept at 4°C until imaging. Nuclei were stained with Hoechst 33342 for 1 h in the dark and cells were imaged using an OperaPhenix confocal microscope with 40 x magnification, 405/456 nm and 640/706 nm, excitation/emission respectively.

### Data presentation and statistical analysis

All statistical analyses were carried out using three independent biological replicates and two or three technical repeats, error bars are standard error of the mean (SEM) and EC50 was interpolated using non-linear regression (inhibitor vs. response-four parameters) in GraphPad Prism Software 8.4.2. Each statistical test was performed on raw data normalised to cell number or signal before normalisation to the percentage of control was performed. Oneway ANOVA with Dunnett’s test comparison to control untreated cells was utilised to calculate significance. P values are indicated in the respective figure legends with symbols (* P < .05 and ** P < .01; **** P < .0001).

### Key resources table

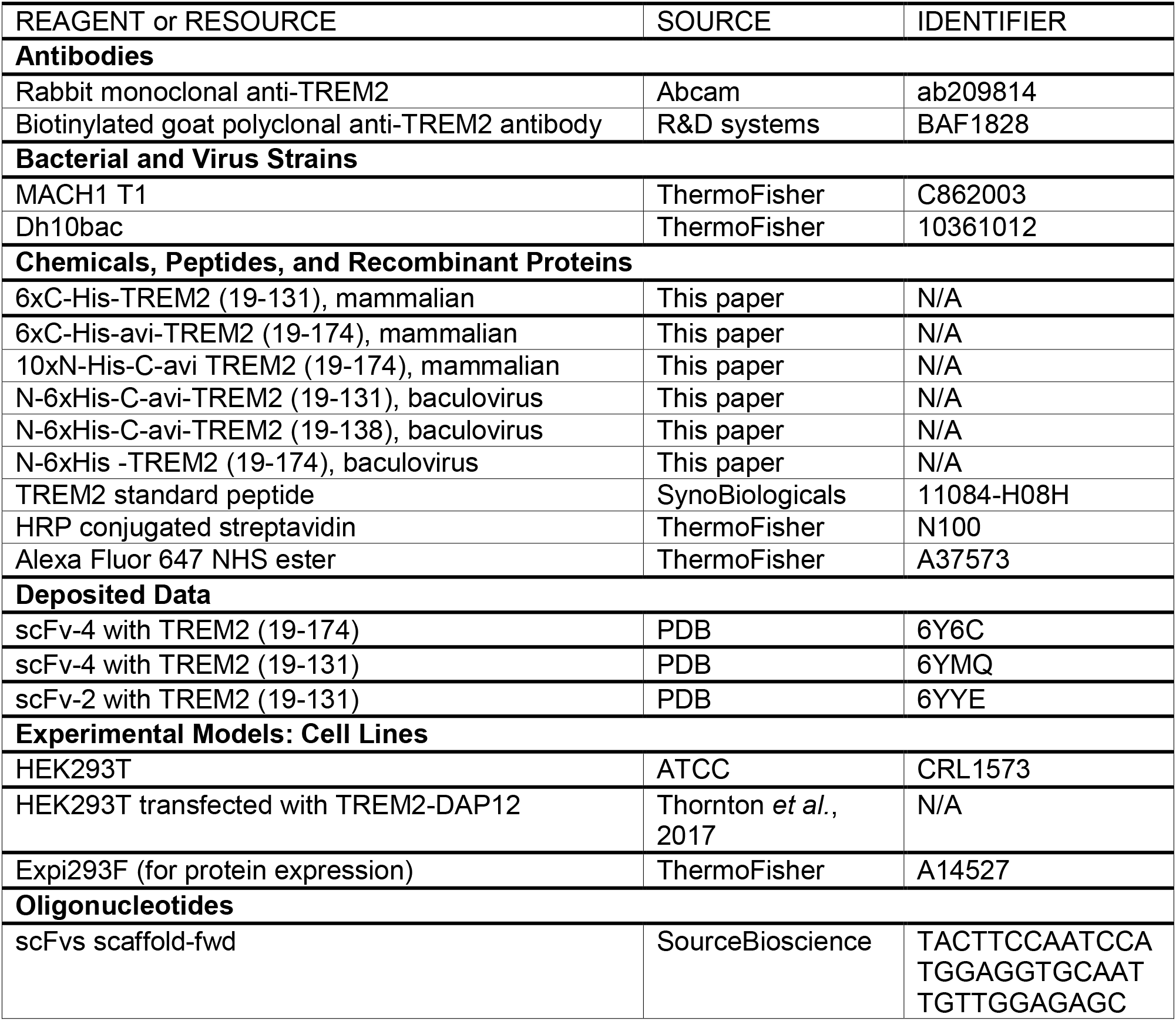
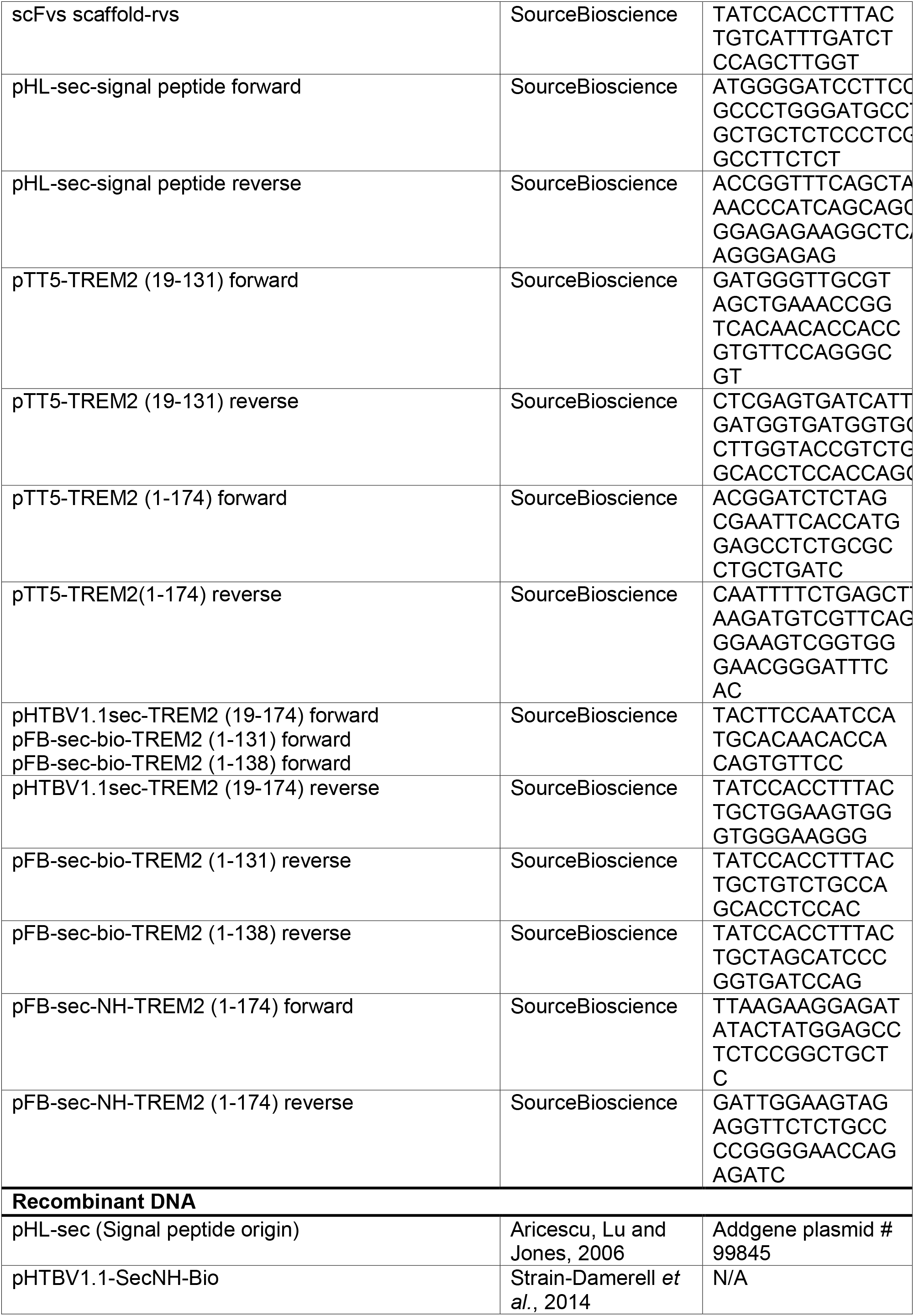
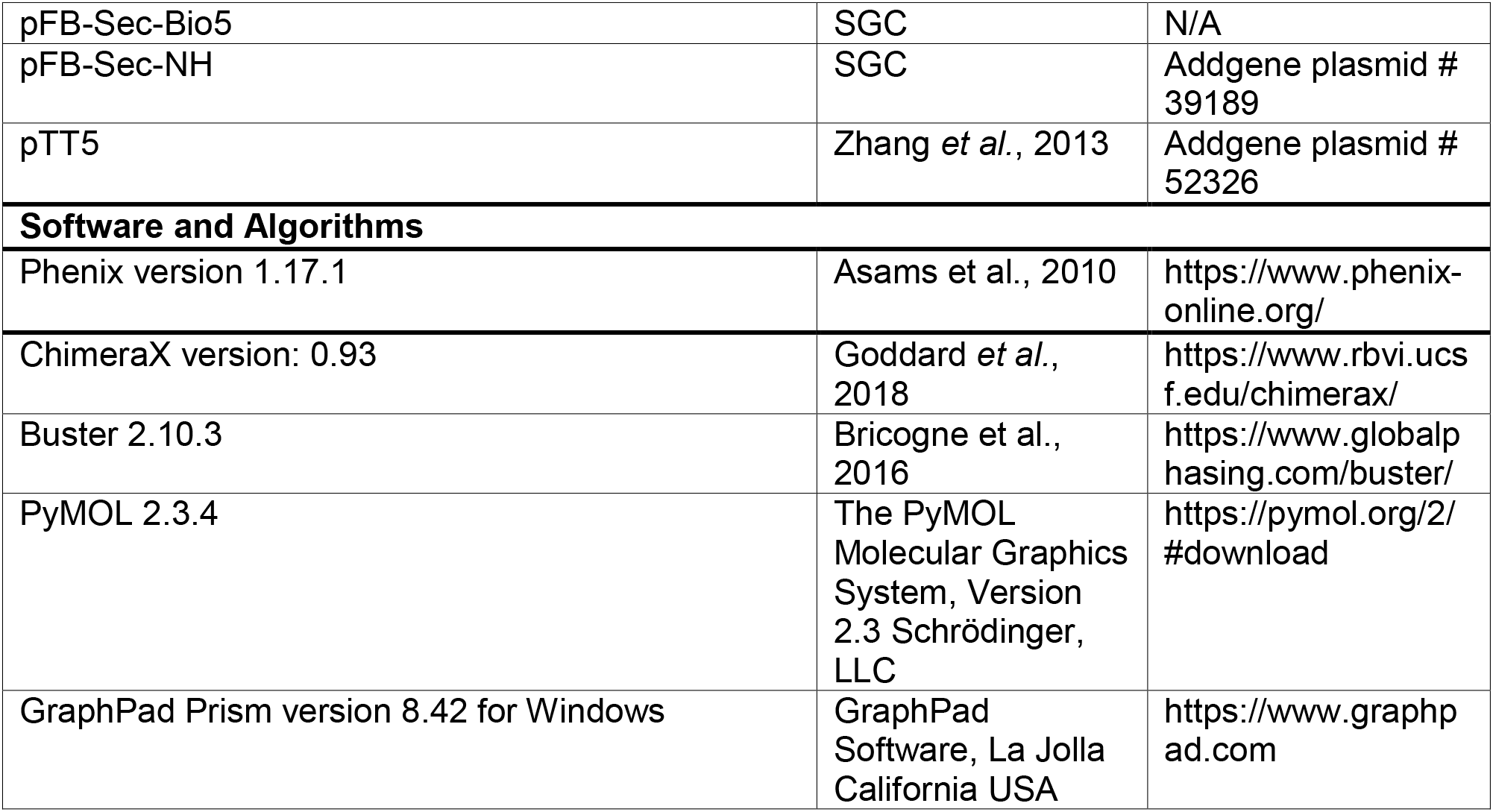

## Notes

### Competing Interest Statement

The authors have declared no competing interest.

