## Supplemental Figures 1-5 for "Selection and structural characterisation of anti-TREM2 scFvs that reduce levels of shed ectodomain"

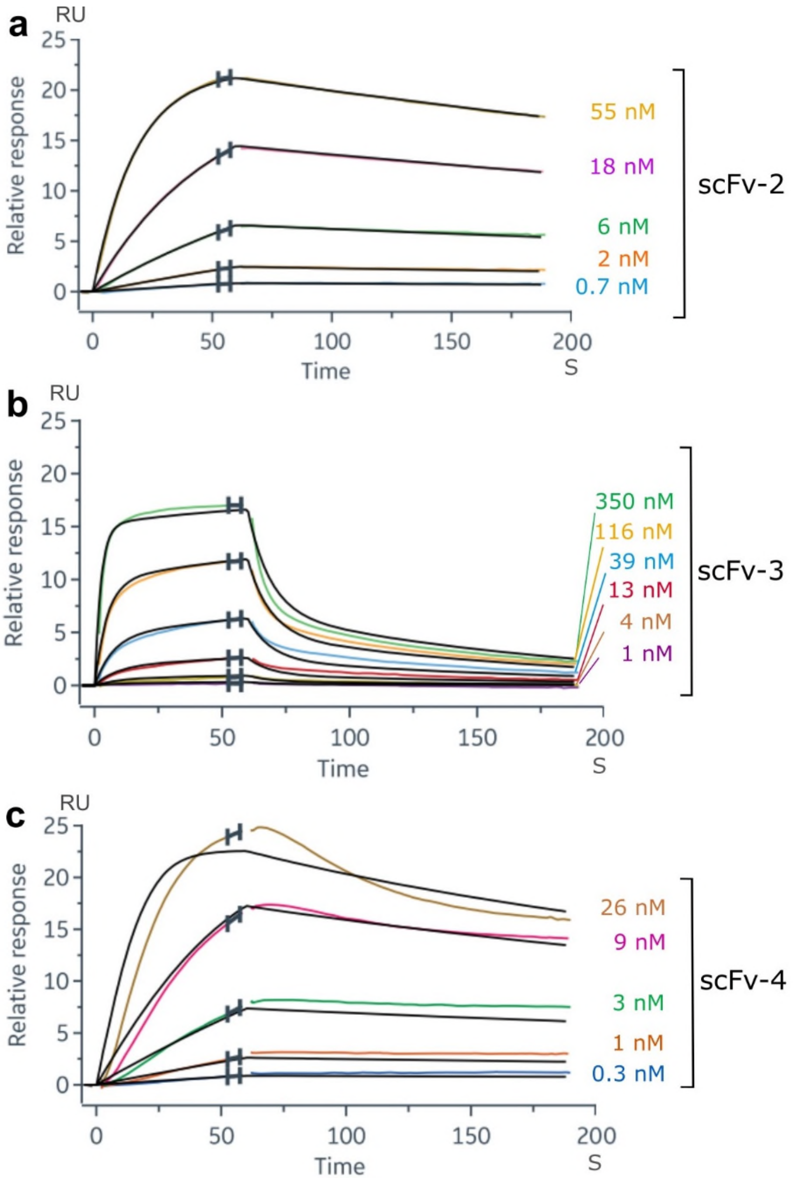


**Figure S1. Kinetic measurements of three anti-TREM2 scFvs and estimation of apparent $K_D$.** Binding kinetics were determined by surface plasmon resonance (SPR) performed on a Biacore 8K instrument. Biotinylated TREM2 (His19-Ser174) produced in mammalian cells was immobilised and binding of different concentrations of scFv containing mixture of monomer and dimer was analysed. (**a,b,c**) Binding of scFv-2 and scFv-4 to TREM2 fitted using a Langmuirian 1:1 model, scFv-3 fit was improved by use of 2:1 binding model. ScFv-2: $k_{on}$ = 9.9 x $10^5$ $M^{-1}s^{-1}$, $k_{off}$ = 1.6 x $10^{-3}$ $s^{-1}$, ${}^{App}K_D$ = 1.6 nM; ScFv-3: $k_{on}$ = 5.0 x $10^5$ $M^{-1}s^{-1}$, $k_{off}$ = 2.7 x $10^{-2}$ $s^{-1}$, ${}^{App}K_D$ = 54 nM; ScFv-4: $k_{on}$ = 5.0 x $10^6$ $M^{-1}s^{-1}$, $k_{off}$ = 2.9 x $10^{-3}$ $s^{-1}$, ${}^{App}K_D$ = 0.6 nM. Some non-specific interaction to reference chip was observed for scFv-3 and scFv-4. The $k_{on}$ and $k_{off}$ may contain avidity component, especially in the case of scFv-3 and scFv-4 so ${}^{App}K_D$ is used to describe the results. Association and dissociation curves are coloured for each concentration and the fitted curves are drawn in black.

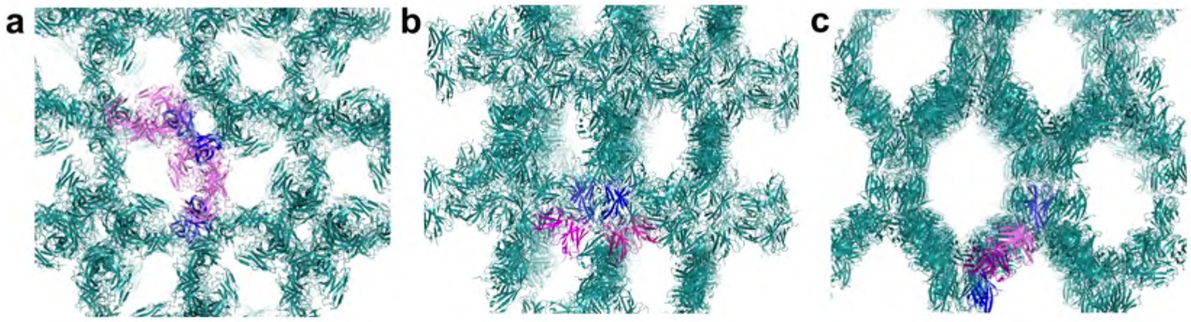


**Figure S2. The crystal lattices of scFvs and TREM2 complexes.** (**a**) Crystal lattice of the scFv-4 and TREM2 (6YMQ) complex crystallised in unit cell of 114, 126, 225 Å and 69% solvent composition. Green lattice visualised by generating symmetry mates of components of asymmetric unit consisting of 6 subunits of scFv (pink) and six subunits of TREM2 (blue) (**b**) Crystal lattice of scFv-4 and TREM2 complex (6Y6C) crystallised in unit cell of 112, 112, 232 Å and solvent composition 74%. Lattice was visualised by generating symmetry mates of components of asymmetric unit consisting of 2 subunits of scFv (pink) and two subunits of TREM2 (blue). (**c**) Crystal lattice of scFv-2 and TREM2 complex (6YYE) crystallised in unit cell of 112, 112, 232 Å and solvent composition 77%. Lattice was visualised by generating symmetry mates of components of asymmetric unit consisting of 2 subunits of scFv (pink) and six subunits of TREM2 (blue).

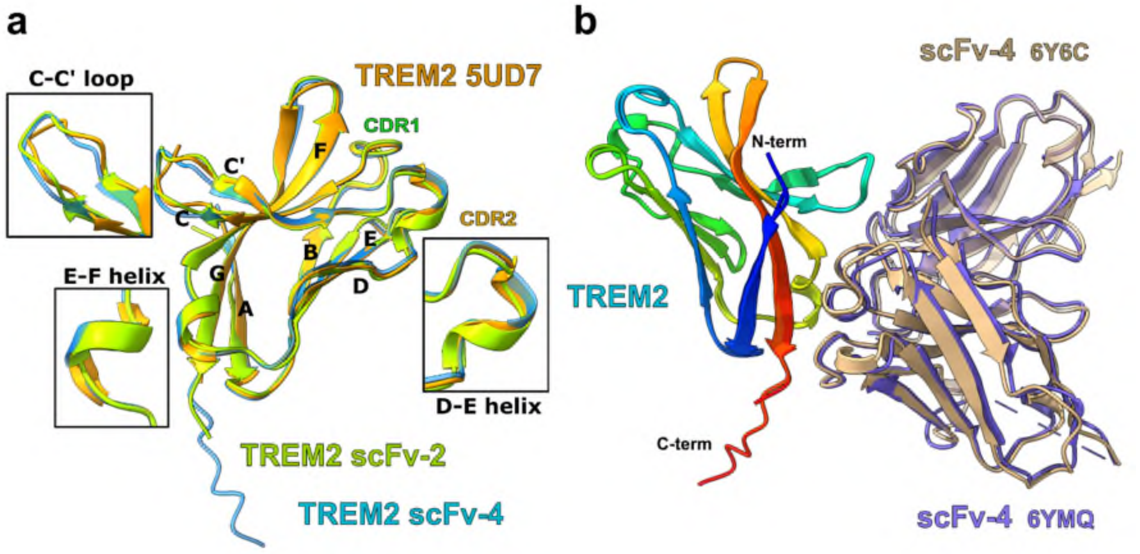


**Figure S3. Superimposition of selected TREM2 and scFv-4 co-structures**. **(a)** Superimposition of the highest resolution structure of TREM2 (2.2 Å) previously deposited in the PDB, 5UD7 (Sudom *et al.*, 2018), and TREM2 bound to scFv-2 and scFv-4. The root mean square deviation (RMSD) of C-alpha atoms comparing 5UD7 to the TREM2 complexes with scFv-2 and scFv-4 is 0.7 Å and 0.4 Å, respectively. The loops which show the highest variation are shown in inset panels and named by secondary structure or TREM2 CDR. Amongst them is the C-C' loop which is differentially bound by the scFvs and the CDR2 loop which shows a variable extent of a short α-helix, especially when bound to scFv-2. TREM2 5UD7 is coloured in orange, scFv-4-bound TREM2 in blue and scFv-2-bound TREM2 in green. **(b)** Superposition of the two TREM2-scFv-4 structures showed excellent agreement (Cα RMSD = 0.5 Å). ScFv-4 in complex with long TREM2 (His19-Ser174, 6Y6C) is shown in tan, whereas scFv-4 in complex with short TREM2 (His19-Asp131, 6YQM) is shown in purple.

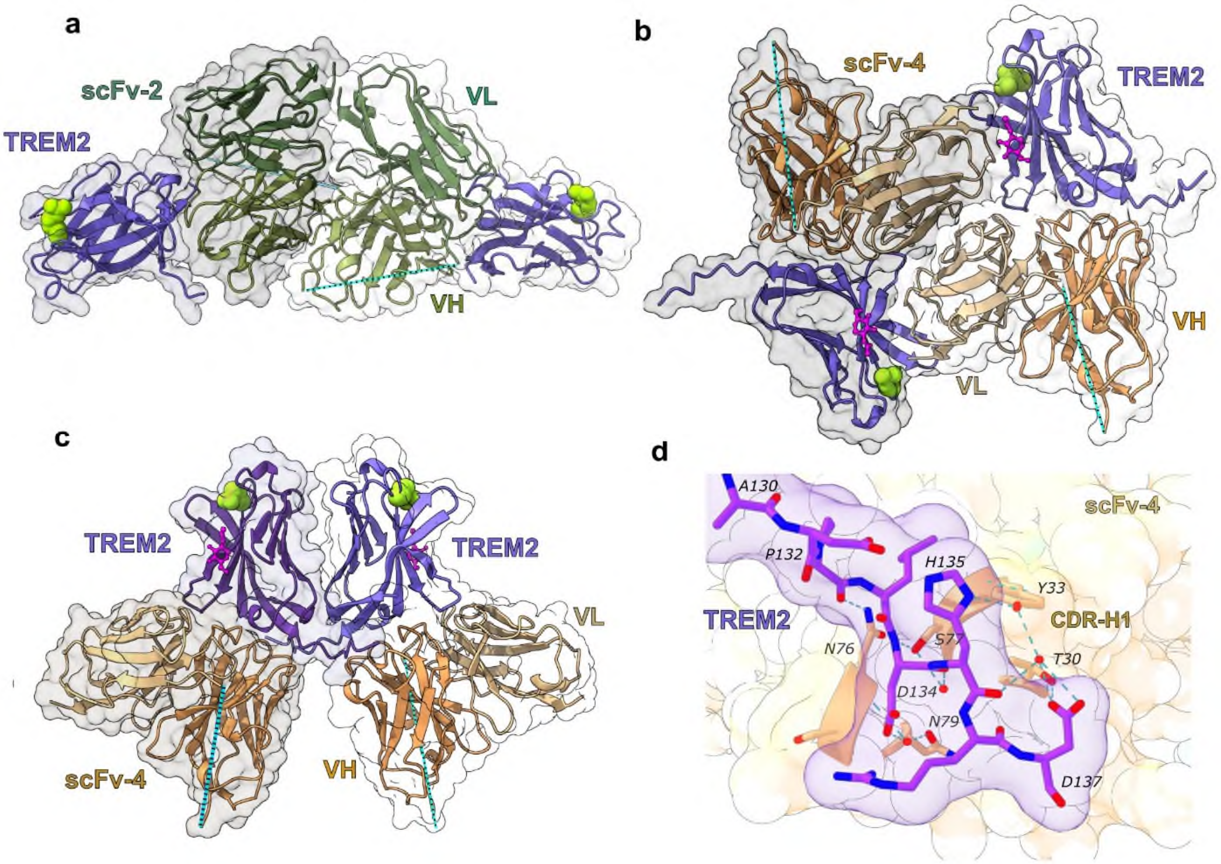


**Figure S4. Different crystal packing of TREM2-scFv4 complexes and scFv-4-stalk interactions.** (**a**) Crystal packing between scFv-2 subunits in the TREM2 complex (PDB 6YYE). The scFv-2 VH and VL domains are coloured in different shades of green; The disordered linker between VH and VL domains is shown by a dotted cyan line. TREM2 is coloured slate blue. A molecular surface representation of each TREM2-scFv-2 complex is shown in a different transparent shade of grey. (**b**) Crystal packing between scFv-4 subunits (tan) observed in the TREM2 (a.a. 174) co-structure (PDB 6Y6C). Each scFv-4 subunit packs against two molecules of TREM2 by interaction with the primary TREM2 epitope as well as an additional surface consisting of the TREM2 C-C' and F-G loops. (**c**) Additional, crystal packing of TREM2 dimer with two scFv-4 observed in the same structure but containing stalk interaction. Each TREM2 subunit interacts with two scFv-4 molecules through interactions from the TREM2 ectodomain and stalk region, respectively. N-acetylglucosamine (NAG) is coloured in pink and presented as ball and stick (**d**) The TREM2 C-terminal stalk sequence Pro132-Asp137 (purple) packs against ScFv-4 CDR-H1 and Asn76-Asn79 (tan). Residues forming hydrogen bond interactions are presented as sticks and waters as red spheres.

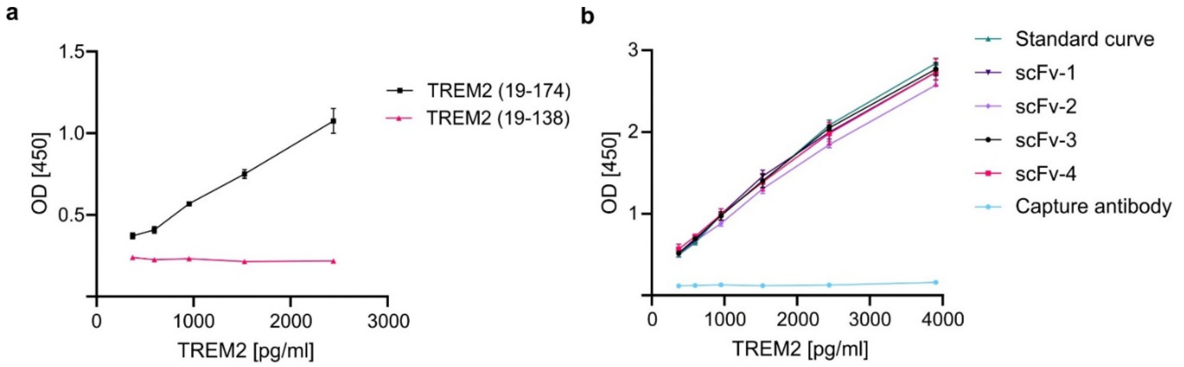


**Figure S5. Validation of the sTREM2 ELISA assay.** (**a**) Identification of the soluble TREM2 amino acid sequence essential for ELISA detection. ELISA was performed using TREM2 (a.a. 19-174) protein containing the immunoglobulin domain and stalk and a shorter immunoglobulin only TREM2 protein (a.a. 19-138) in order to determine the requirement for the stalk sequence. The data represent n=1, 2 technical repeats. (**b**) Characterisation of epitope compatibility between ELISA antibodies and scFvs. Each of the four scFvs was incubated at 10 µg/mL with 3906 pg/mL standard curve peptide for 15 min. Dilution series were used to generate curves and a standard curve containing TREM2 peptide only was used for comparison. The capture antibody added to the standard curve is used to demonstrate the effects of overlapping epitopes. The data represent n=1, 2 technical repeats, except for scFv-3 and scFv-4 where n=2 independent experiments.
